# Laboratory evolution and polyploid SCRaMbLE reveal genomic plasticity to synthetic chromosome defects and rearrangements

**DOI:** 10.1101/2022.07.22.501046

**Authors:** Thomas C. Williams, Heinrich Kroukamp, Xin Xu, Elizabeth L. I. Wightman, Briardo Llorente, Anthony R. Borneman, Alexander C. Carpenter, Niel Van Wyk, Monica I. Espinosa, Elizabeth L. Daniel, Roy S. K. Walker, Yizhi Cai, Helena K. M. Nevalainen, Natalie C. Curach, Ira W. Deveson, Timothy R. Mercer, Daniel L. Johnson, Leslie A. Mitchell, Joel S. Bader, Giovanni Stracquadanio, Jef D. Boeke, Hugh D. Goold, Isak S. Pretorius, Ian T. Paulsen

**Author notes:** Corresponding authors: Thomas C. Williams,. Isak S. Pretorius,. Ian T. Paulsen. Present affiliations (Roy S. K. Walker, Yizhi Cai).

## Abstract

We have designed, constructed, and debugged a synthetic 753,096 bp version of *Saccharomyces cerevisiae* chromosome XIV as part of the international Sc2.0 project. We showed that certain synthetic loxPsym recombination sites can interfere with mitochondrial protein localization, that the deletion of one intron (*NOG2*) reduced fitness, and that a reassigned stop codon can lead to a growth defect. In parallel to these rational debugging modifications, we used Adaptive Laboratory Evolution to generate a general growth defect suppressor rearrangement in the form of increased *TAR1* copy number. We also extended the utility of the Synthetic Chromosome Recombination and Modification by LoxP-mediated Evolution (SCRaMbLE) system by engineering synthetic-wild-type tetraploid hybrid strains that buffer against essential gene loss. The presence of wild-type chromosomes in the hybrid tetraploids increased post-SCRaMbLE viability and heterologous DNA integration, highlighting the plasticity of the *S. cerevisiae* genome in the presence of rational and non-rational modifications.

## Introduction

The field of synthetic genomics encompasses the design, construction, and characterization of whole genomes and chromosomes. This new approach to genomics provides several unique opportunities. For example, the ability to make global genetic changes that are too numerous to implement in a step-wise manner, the capacity to discover new biological phenomena through the classic ‘design-build-test-learn’ cycle of synthetic biology, and the potential to design genomes that encode superior industrial phenotypes (Pretorius, 2017; Pretorius and Boeke, 2018) are all enabled by synthetic genomics. The synthetic *Saccharomyces cerevisiae* genome project ‘Sc2.0’ exemplifies these new possibilities via genome streamlining (removal of transposons and non-essential introns), genome ‘defragmentation/refactoring’ via the relocation of all transfer RNA genes to a separate neo-chromosome, telomere standardisation, and through the placement of heterologous loxPsym recombination motifs just after the stop codon of every non-essential gene. The 12 Mb *S. cerevisiae* genome consists of sixteen chromosomes, built by an international consortium adhering to central design principles (Richardson et al., 2017). The Sc2.0 consortium has already described six and one half synthetic yeast chromosomes, resulting in new fundamental biological knowledge and genome construction technology. For example, novel growth defect ‘debugging’ (Wu et al., 2017) and chromosome consolidation (Mitchell et al., 2017) techniques have been developed, an in-depth phenotypic characterization of designed chromosomes (Shen et al., 2017; Xie et al., 2017) and the degree of genome plasticity with regard to ribosomal gene clusters (Zhang et al., 2017), and the effects of chromosome re-design on the three-dimensional genomic architecture have been investigated (Mercy et al., 2017).

The most significant design feature incorporated in Sc2.0 is an inducible evolution system termed SCRaMbLE (Synthetic Chromosome Recombination and Modification by LoxP-mediated Evolution). Induction of a heterologous Cre-recombinase enzyme results in inversions, duplications, translocations, and deletions of genes between LoxP sites (Annaluru et al., 2014; Dymond and Boeke, 2012; Dymond et al., 2011; Jovicevic et al., 2014; Mercy *et al*., 2017; Shen et al., 2016). The induction of SCRaMbLE can in theory generate a virtually unlimited number of genomes with unique gene content and genomic architecture (Shen *et al*., 2016), making it an extremely powerful tool for generating genetic diversity prior to laboratory evolution experiments, and for understanding the genomic basis of selected phenotypes (Vickers et al., 2017; Williams et al., 2016). However, there are significant limitations to SCRaMbLE in its current form. For example, due to the relatively high incidence of gene deletions there is a high frequency of lethal modifications in a SCRaMbLE’d population, significantly reducing genomic diversity. This problem has been partially solved through the use of synthetic-wild-type heterozygous diploid strains, where the presence of non-SCRaMbLE-able chromosomes buffers against essential gene loss (Jia et al., 2018; Shen et al., 2018). In addition to issues with lethality, SCRaMbLE is predominantly used to vary the gene content of synthetic yeast chromosomes, pathway-encoding linear DNA, or plasmids (Liu et al., 2018). An ideal scenario would be for SCRaMbLE to give rise to the highest possible level of genetic variation without excessive cell death, and simultaneously enable the incorporation of multiple heterologous gene expression cassettes.

In addition to the construction and debugging of synthetic chromosome XIV (synXIV), we have improved the SCRaMbLE system by varying the number of synthetic and wild-type chromosomes in a series of novel hybrid tetraploid strains, and functionalizing heterologous gene expression cassettes with loxPsym recombination sites for post-transformation SCRaMbLE in these strains.

## Results

### SynXIV design and construction

*S. cerevisiae* synXIV was redesigned according to Sc2.0 principles using the BioStudio software package (Richardson *et al*., 2017). Briefly, 256 loxPsym sites were inserted 3 bp after the stop codons of non-essential genes, 14 introns were removed, ORFs were synonymously recoded to contain a total of 1040 PCR-tags, 90 stop codons were swapped to ‘TAA’ to free-up the TAG codon for potential future reassignment, native telomeres were replaced with standardized synthetic versions, and all transposon and tRNA sequences were removed. These changes resulted in a 753,097 bp synXIV divided into 24 ‘megachunks’ labelled A to X (File S1), representing a 4% size reduction of the native 784,333 bp version. SynXIV was constructed according to the Sc2.0 Swap-In approach (Richardson *et al*., 2017) across two different strains that were crossed to generate a near complete version of synXIV (Figure S1A).

### SynXIV characterization and debugging

#### LoxPsym insertion in the 3’ UTR of *MRPL19* causes a respiratory growth defect

The synXIV strain with megachunks (i.e., ∼30-60 kb synthetic DNA fragments) G-X showed a growth-defect after the integration of the first megachunk, which was shared with all subsequent strains. However, the growth defect was not observed when megachunk G was re-integrated in a wild-type BY4741 strain. Whole-genome re-sequencing of the original megachunk G integrant strain revealed that chunks (i.e., ∼10 kb synthetic DNA fragments) G1 and G2 had been integrated approximately four times, as indicated by coverage relative to surrounding chromosomal loci. When a descendant of this strain with megachunks G-O integrated was sequenced, the multiple copies were no longer present, and we conclude it had presumably been spontaneously looped out of the genome via homologous recombination. However, this strain and all subsequent megachunk integration strains shared a severe growth defect on YP-glycerol medium. To ascertain the cause of this problem, backcrossing and pooled fast/slow strain sequencing was carried out as previously described (Wu *et al*., 2017). A synthetic chromosome region spanning megachunks J to L was found to have low-coverage in ‘fast-grower’ pool reads. Conversely, a wild-type chromosome region from megachunks H to L had low-coverage in the pooled ‘slow-grower’ reads (Figure S1 C, D), suggesting that a synXIV modification within megachunks J to L led to the defect.

An independent line of inquiry also indicated that the megachunk J to L region was the cause of a major growth-defect, and further narrowed the location down to chunk J1. During the final meiotic cross of partially synthetic strains to produce a fully synthetic version of chromosome XIV (Figure S1B), two near-complete strains were identified that had wild-type I-J and J regions, respectively. These strains had improved fitness relative to two strains with fully-synthetic versions of chromosome XIV (Figure S1B), suggesting that the cause of a growth defect lay within the megachunk I-J region, independently supporting the back-crossing and pooled sequencing analysis. Integration of megachunk I in one of these faster-growing strains (SynXIV-29) did not cause any growth defect, indicating that the defect lay outside megachunk I. When synthetic chunk J1 was then introduced, the severe growth defect on YP-glycerol was re-established (Figure 1A). Subsequent integration of the wild-type J1 region did not restore normal growth, initially leading us to dismiss this region as the cause of the growth defect. During the integration of synthetic chunk J1, several strains were identified as having correct integration according to PCR-tag analysis, and one of these strains (J1.8) was found not to have the growth defect (Figure 1A). Whole genome sequencing of slow and fast-growing versions of the J1 integrants revealed that the fast growing isolate J1.8 was missing a single loxPsym site immediately 3′ of the *MRPL19* gene (Figure 1B), whereas in the slow-growing isolate (J1.4), this loxPsym was present. The *MRPL19* gene encodes a mitochondrial ribosomal protein, and deletion of this gene causes a respiratory growth defect (Merz and Westermann, 2009). Further analysis of the re-sequenced genomes showed that the slow-growing isolate had no reads mapping to the yeast mitochondrial reference genome, while the fast-growing isolate did. Loss of mitochondrial DNA is consistent with the fact that re-integration of wild-type chunk J1 did not restore growth, as yeast cannot de-novo regenerate the mitochondrial genome once it has been lost (Parisi et al., 1993). It is also consistent with the complete lack of respiratory growth on YP-glycerol seen from this defect (Figure 1A).

**Figure 1.**
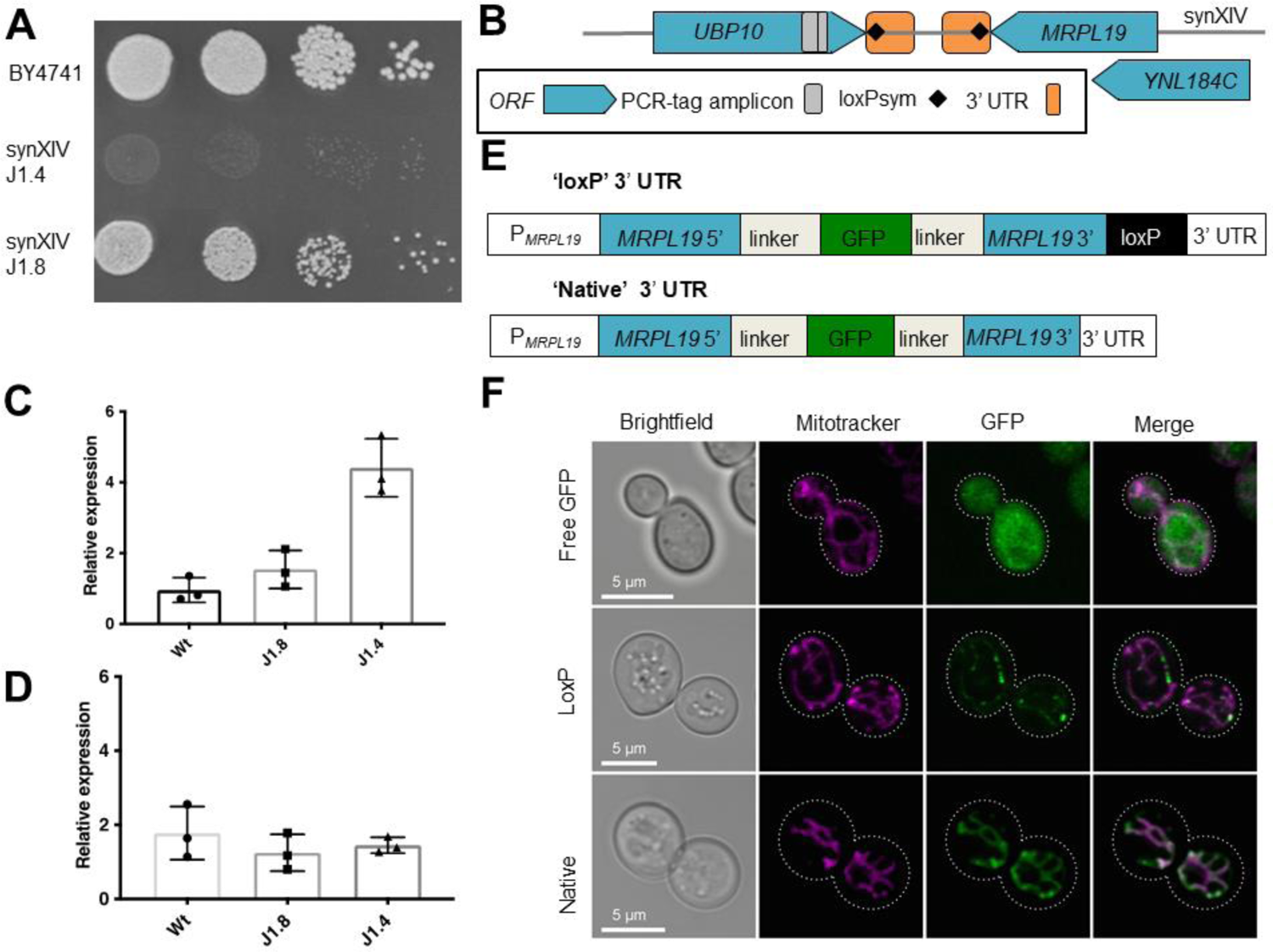
Chunk J1 growth defect and gene expression analysis. (**A**) YP-glycerol fitness test of chunk J1 integrants 4 and 8 (strains 39 and 40, Table S1) and the Wt (BY4741, strain 1, Table S1). The plate was incubated at 30°C for 4 d prior to imaging, and is representative of two repeated experiments. (**B**) Genetic context of the *MRPL19* gene and the surrounding synthetic chromosome design features. (**C**) RT-qPCR of the *MRPL19* and (**D**) *NPR1* genes was carried out on cDNA from BY4741 (Wt), repaired synXIV (J1.8, strain 40, Table 3), and growth defect synXIV (J1.4, strain 39, Table S1) strains. Expression was normalised to the *ALG9* gene using the modified-Livak method as previously described (Williams et al., 2015). Bars and error bars represent mean and standard deviation from three biological replicates. Individual expression values of replicates are also shown. (**E**) Two synthetic *MRPL19* promoter-gene-3’UTR constructs were designed with a super folder GFP expressed from the middle of the native ORF, separated by peptide linkers. One version contained a loxPsym motif 3 bp after the stop codon (termed ‘loxP’) while the second version contained no loxP within the native 3’UTR (termed ‘Native’). (**F**) BY4741 strains expressing either of these two constructs (strains 49 and 48), or a cytosol localized GFP (termed ‘Free GFP’ (Williams et al., 2017), strain 50, Table S1) were grown in the presence of 100 nM Mitotracker Red (ThermoFisher) to stain mitochondria. An Olympus FV 1000 confocal microscope was used to visualize yeast cells with bright field, mitotracker and GFP signals. See also Figure S1.

We hypothesized that the presence of a loxPsym site in the 3′ UTR of *MRPL19* could modulate transcriptional termination efficiency and hence performed RT-qPCR on RNA extracted from exponentially growing wild-type, J1.4, and J1.8 strains to test this. Interestingly, *MRPL19* transcript levels were significantly up-regulated by approximately 5-fold in the slow-growing J1.4 isolate, but were not significantly different between the wild-type and fast-growing J1.8 isolate (Figure 1C, 2-sided student t-test, p < 0.05). The mRNA levels of the nearby *YNL184C* and *NPR1* ORFs were also measured to determine if the *MRPL19* 3′ UTR loxPsym site had any effects on their transcript levels. No mRNA was detected from the *YNL184C* ORF in any of the strains, while *NPR1* expression levels were not significantly different between the strains (Figure 1D). To test whether the observed up-regulation of *MRPL19* mRNA could cause the growth defect in the slow-growing J1.4 strain, the native *MRPL19* gene and terminator were over-expressed from the strong-constitutive *TDH3* promoter (Peng et al., 2015) in the wild-type strain from the pRS413 vector. *MRPL19* over-expression did not cause a growth defect, suggesting the mechanism of the growth defect is unrelated to *MRPL19* over-expression.

The *MRPL19* mRNA has a Puf3p recognition motif, and when *PUF3* is deleted there is no *MRPL19* mRNA localization to the mitochondria (Saint-Georges et al., 2008). The addition of a loxPsym site to the 3′ UTR of *MRPL19* might therefore interfere with mitochondrial mRNA targeting, leading to the observed growth defect. To test this hypothesis, we designed a GFP fusion protein that retained the entire *MRPL19* coding sequence in order to account for the possibility that other RNA or protein signals are important for mitochondrial protein import (Figure 1E). Versions with the native 3′ UTR and with the loxPsym containing 3′ UTR were synthesized and tested for the import of GFP into yeast mitochondria (Figure 1F). Confocal microscopy of yeast cells with stained mitochondria (mitotracker) showed that the cytosol-localized control (‘Free GFP’) had no GFP signal correlation with the mitotracker signal, while the native *MRPL19-GFP* construct (‘Native’) resulted in co-localization of GFP with the mitochondria (Figure 1F). In contrast, the insertion of a loxPsym motif in the 3′ UTR of *MRPL19* appeared to interfere with mitochondrial localization and import, as GFP signal was commonly clustered around mitochondria (Figure 1F and Figure S1 E, F). Taken together with the observed growth defect, it is therefore likely that the loxPsym motif in the 3′ UTR of *MRPL19* interferes with correct mitochondrial localization of this protein, leading to the observed growth defect under respiratory conditions (Figure 1A).

### Adaptive Laboratory Evolution restores respiratory growth through increased *TAR1* copy number

Although the removal of the *MRPL19* 3′ UTR loxPsym dramatically improved growth on both YPD and YP-glycerol, there was still an obvious growth defect on YP-glycerol medium at 30°C (Figure 1A). To repair and understand the cause of this defect, the J1.8 strain was subjected to Adaptive Laboratory Evolution (ALE) in liquid YP-glycerol medium in triplicate for approximately 90 generations by passaging into fresh medium every 24 h (Figure 2A). The wild-type BY4741 strain was also evolved in parallel to enable the exclusion of mutations that enhance glycerol utilization, or mutations related to general adaptation to YP media, and to assess the accumulation of ‘hitch-hiker’ mutations occurring due to genetic drift under these conditions. Initially, each passage was inoculated at an A_600_ of 0.1, but inoculation was decreased to 0.05 after 24 generations to enable faster accrual of generations and DNA replication errors. A_600_ was measured after each 24 h period to serve as a proxy for fitness. The J1.8 strain showed a 50 % improvement in final A_600_ on YP-glycerol medium after 90 generations, whereas the wild-type control strain showed only a 38 % improvement after 120 generations (Figure S2). Fitness testing of the wild-type strain (BY4741), parental synXIV strain (J1.8), and a mixed population from one of the J1.8 evolutionary lineages (J1.8e3i) revealed that growth on YP-glycerol was restored to wild-type levels (Figure 2B).

**Figure 2.**
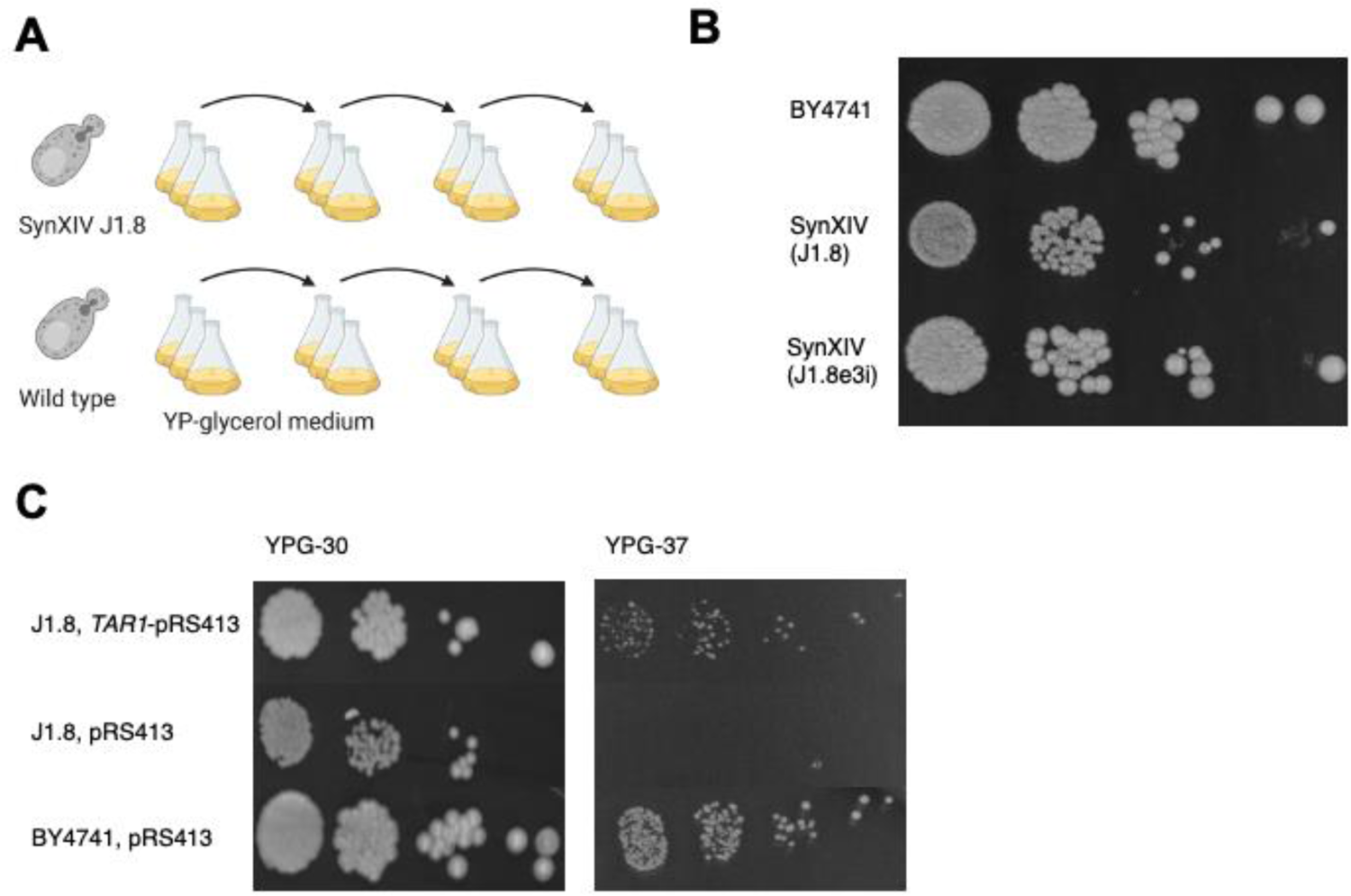
Adaptive laboratory evolution of synXIV and Wt strains on YP-glycerol medium. (**A**) BY4741 (Wt, strain 1, Table S1) and synXIV strains (J1.8, strain 40, Table S1) were grown in YP-glycerol medium with passaging to fresh medium every 24 h. (**B**) At the end of the evolution experiment the fitness of the parental wild-type and J1.8 strains were compared with one of the evolved J1.8 lineages (J1.8e3i, strain 47, Table 3) on YP-glycerol at 30°C. (**c**) Fitness test of the SynXIV intermediate strain J1.8 (strain 40, Table S1) with and without *TAR1* expression from its native promoter on the pRS413 plasmid in YP-glycerol at 30 and 37°C. BY4741 (Wt, strain 1, Table S1) transformed with empty pRS413 plasmid is shown as a control. Photos were taken after 5 d and are representative of repeated experiments. The image in panel A was made using Biorender.com. Related to Figure S2.

Whole-genome sequencing of isolates and final evolved mixed populations was carried out to compare mutations that might have caused the initial growth defect during the construction of synXIV. While no point mutations were detected anywhere in the genomes of the J1.8 evolved lineages that were absent from the control lineages, they did have a higher relative copy number of the ribosomal DNA repeat located on chromosome XII, with the evolved J1.8 lineages sharing approximately eight more rDNA copies compared to the parental J1.8 strain. The *TAR1* gene is encoded antisense to the *RDN25-1* gene on the rDNA locus, and plays a role in the quality control of defective mitochondria (Bonawitz et al., 2008), particularly when mixed populations of defective and functional mitochondrial populations are inherited after mating (Poole et al., 2012). The Tar1p response to defective mitochondria is mediated via the formation of extra-chromosomal rDNA circles (ERCs) which relieve *TAR1* expression from Sir2p mediated repression by physically locating the gene away from the native chromosomal locus (Li et al., 2006; Poole *et al*., 2012). This process occurs as a result of the yeast retrograde response, which facilitates glutamate synthesis in the absence of a complete TCA cycle in defective mitochondria (Jazwinski and Kriete, 2012). Long-read nanopore sequencing and *de-novo* assembly of evolved J1.8 isolate genomes not only resulted in full-length contigs for each of the 16 chromosomes (File S2), but also in additional contigs containing *TAR1*-rDNA repeats. These extra rDNA repeats were not observed in the genome sequences of evolved wild-type populations and were not contiguous with the chromosome XII sequence, suggesting that they represent extra *TAR1* copies presumably in the form of ERCs in the J1.8 evolved isolates. This phenomena of circular extrachromosomal *TAR1*-encoding circular DNA has previously been observed in yeast cells with defective mitochondria (Borghouts et al., 2004). Expression of the *TAR1* gene from its native promoter on a pRS413 vector improved growth of the parental synXIV strain on YP-glycerol at 30°C and 37°C (Figure 2C), suggesting that increased rDNA copy-number enabled higher *TAR1* expression and normal respiratory growth in the evolved lineages.

### SynXIV-wild-type backcrossing restores synXIV fitness

Ectopic expression of *TAR1* in synXIV was used to improve growth based on knowledge gained from ALE (Figure 2). However, this solution was not satisfactory for the completion of synXIV debugging for two reasons. Fitness was not fully restored to wild-type levels, and it is possible that *TAR1* expression simply suppresses defects caused by either synXIV design changes or background genomic mutations. If synXIV bugs existed, then it was important that they be fixed to enable normal growth of the final consolidated Sc 2.0 strain without a requirement for *TAR1* over-expression. If genomic mutations that were acquired during the strain construction process were causing the observed discrepancy in fitness between synXIV (J1.8) and the wild-type, then it is important that they be corrected so that the synthetic chromosome consolidation process is not negatively affected. To investigate potential additional bugs in the synXIV J1.8 strain, it was backcrossed to the wild-type BY4742 strain. Haploid colonies resulting from individual randomly isolated spores were fitness tested. There was a mixture of fast, slow, and intermediate growth phenotypes across these isolates, suggesting that more than one locus was contributing to the slow growth phenotype. Individual fast- and slow-growing spores were whole-genome sequenced, and the synthetic/wild-type complement of chromosome XIV was mapped in each case. In both the fast- and slow-growing spores there were no synthetic or wild-type regions of chromosome XIV that clearly correlated with growth (Figure 3 A and B). Additionally, there was one slow-growing haploid that had a completely wild-type version of chromosome XIV, and two fast-growing spores that had almost complete versions of synXIV (12c and 7c). These observations suggested that there were mutations elsewhere in the genome that contributed to the slow-growth phenotype. Background mutations (outside of synXIV) that were not present in the original wild-type parental strain that were present in the backcrossed isolates included *MSH1^P80A^*, *ATP1^A424T^*, *ATP3^I303V^, IRA1^A1259D^, CMR1^P87L^, DIA3^Y273F^*, and *PDR5^P496T^*. This complement of genes and strains had two interesting features. Firstly, there was an enrichment of genes associated with mitochondrial processes such as *ATP1*, *MSH1*, and *ATP3*. Secondly, 11/13 fast growing isolates had the mutated *IRA1* gene, while 14/16 slow growing isolates did not, suggesting this mutation might suppress defects encoded by either synXIV genes or other background mutations. Loci that were over-represented in the slow growing isolates and under-represented in the fast-growing isolates included *PDR5* (69 % compared to 31%) and megachunk W (69 % compared to 38 %). In order to remove deleterious background mutations and generate a synXIV strain with wild-type fitness, the fast-growing 12c and 7c (52 and 53, Table S1) strains were crossed and the resulting haploids screened for both fitness and synXIV completeness. One strain was identified (synXIV.17 strain 55, Table S1) that had both wild-type fitness and a near-complete synthetic chromosome XIV, with wild type DNA only present in megachunk D, W, and X regions. All background mutations were absent from synXIV.17 except *IRA1^A1259D^*. Subsequent correction of the *IRA1* mutation on chromosome II with the wild-type sequence in this strain had no effect on fitness (Supplementary Figure 3C, strain 71, Table S1). This backcrossing process successfully generated a partially synthetic version of chromosome XIV in a genetic background free of deleterious mutations on other chromosomes. In addition to removing these putative deleterious background mutations, it was still possible that synthetic DNA in the megachunk D, W, and X regions could cause a growth defect, as these regions were wild-type in the fast-growing synXIV.17 isolate. We therefore re-integrated megachunks D, W, and X, in synXIV.17 while closely monitoring growth phenotypes.

**Figure 3.**
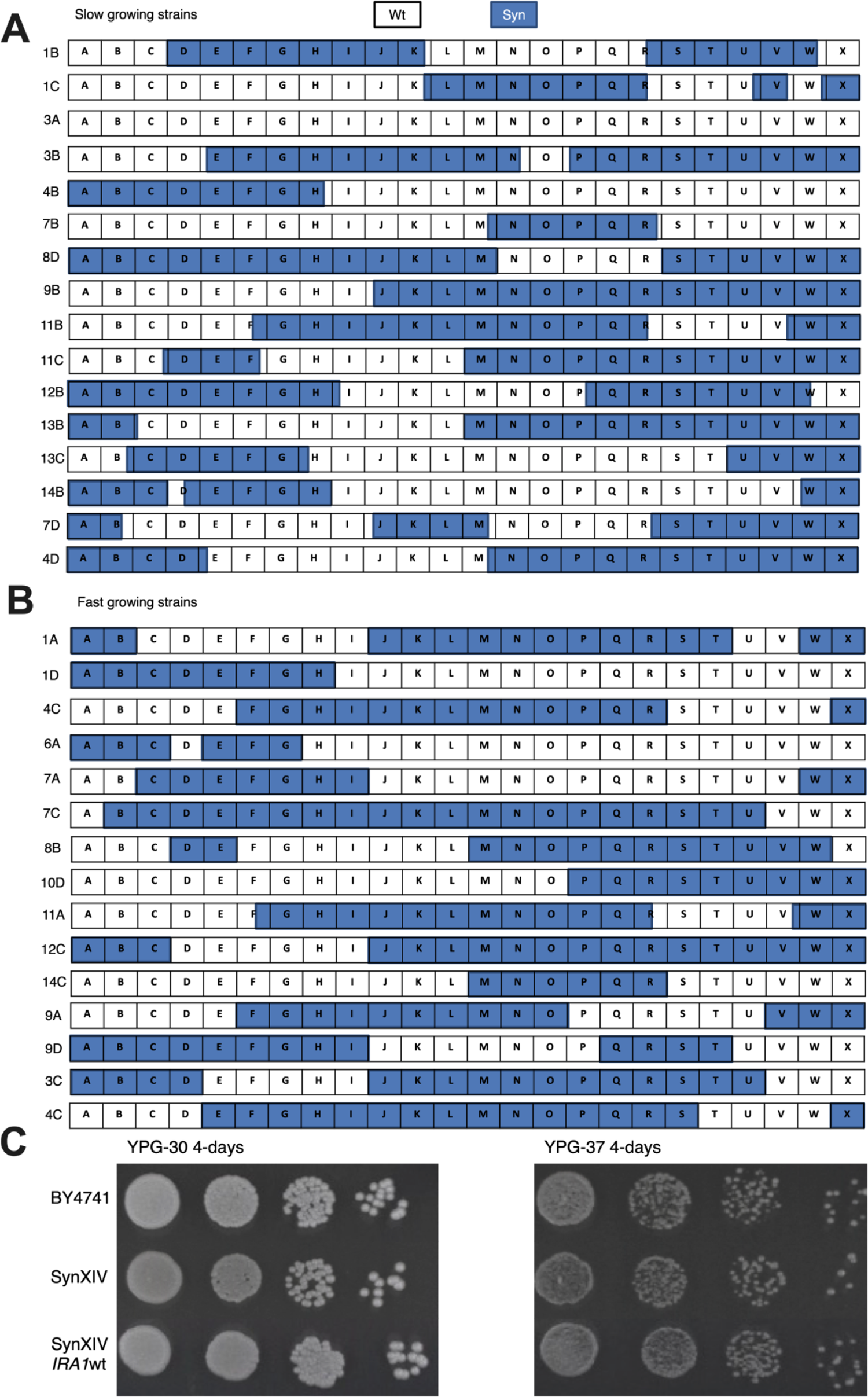
Synthetic DNA regions of chromosome XIV in haploid progeny of a synXIV-BY4741 meiotic cross. (**A**) Slow- and (**B**) fast-growing strains were tested for their synthetic DNA content using PCR tag analysis. Megachunk regions A-X of chromosome XIV are displayed for each strain with blue representing synthetic DNA and white wild-type DNA. All strains are haploid derivates of a cross between strains 1 and 40 (Table S1). (C) Serial 10-fold dilutions of wild-type (BY4741), synXIV (strain 70, Table S1), and SynXIV with the *IRA1^A1259D^* mutation reverted to wild type (strain 71, Table S1). Each strain was plated on YP-glycerol (YPG) at 30 and 37°C for 4 d prior to imaging.

Re-integration of megachunk W in the fast-growing near-complete synXIV.17 led to a fitness defect on YPG at 37°C (strain 57, Table S1), which was absent from a strain with only chunks W3 and W4 present (Figure 4 A, strain 58, Table S1). The main Sc2.0 design change present on chunks W3 and W4 was the removal of the *NOG2* intron, which encodes a small nucleolar RNA (*snr191*) previously shown to cause a growth defect when deleted (Badis et al., 2003). There were no differences in *NOG2* mRNA and protein expression with and without the *snr191* encoding intron present (Figure S3 A-C), while reintroduction of the *NOG2* intron into the synXIV chromosome (Figure 4 B) or via a plasmid (Figure S2 D) restored fitness to wild-type levels. Functional expression of the *NOG2* intron was therefore important for growth independent of Nog2p and *NOG2* mRNA levels (Figure S3), and the intron was retained in synXIV. Similar to the *NOG2* intron, the *SUN4* intron located on megachunk P is also a ‘stable’ intron that accumulates under stress conditions (Morgan et al., 2019). Removal of the *SUN4* intron caused a minor growth defect in the complete synXIV strain (Figure S3E, strain 64, Table S1, strain version number: yeast_chr14_9_01), leading us to retain the intron in the final design (yeast_chr14_9_04).

**Figure 4.**
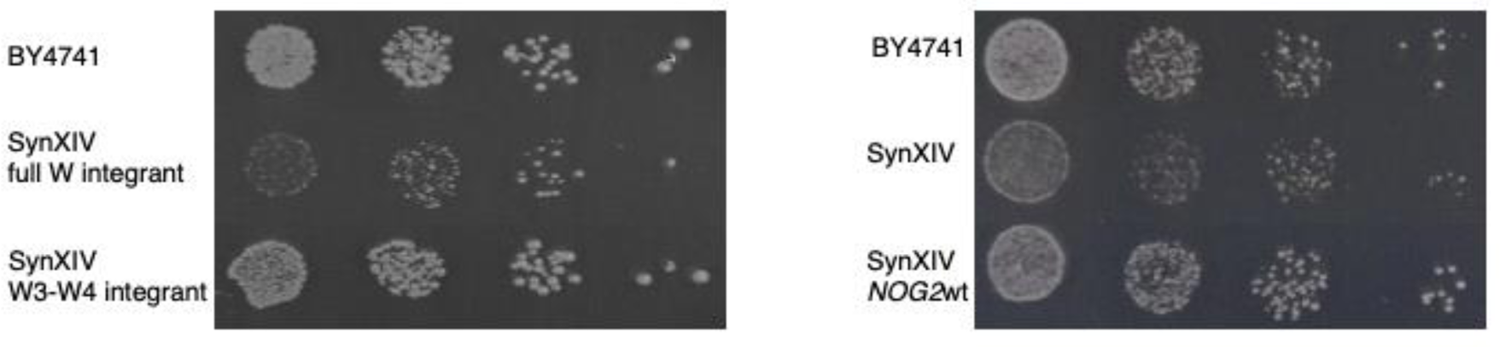
*NOG2* intron growth defect analysis. (**A**) Integration of megachunk W (strain 57, Table S1), but not chunks W3-W4 (strain 58, Table S1) causes a growth defect in SynXIV.17. (**B**) Re-insertion of the *NOG2* intron in the synXIV.17c strain restores wild-type fitness (strain 63, Table S1). Spot assays are 10-fold serial dilutions of exponentially growing cultures on YP-glycerol medium at 37°C. Images were taken after 5 d (**A**) and 3 d (**B**). See also Figure S3.

### Sequence discrepancy repair reveals a growth defect associated with *YNL114C* stop codon reassignment

Sequences that deviated from the intended synXIV sequence were introduced during the construction and debugging of synXIV, with important features repaired to make the final strain (Table S2). While features such as missing PCR tags, restriction enzyme ligation sites, and absence of a small number of LoxP motifs is not critical for the intended function of the synthetic yeast genome, other features, such as stop codon reassignment are expected to be critical in the event of future reassignment of the TAG codon. Additionally, it was possible that non-synonymous mutations in coding sequences could contribute to unidentified phenotypes. All erroneously remaining TAG stop codons were therefore swapped to TAA, and non-synonymous coding sequences were repaired to produce the final synXIV strain (Table S3). During this process we discovered that the introduction of TAA stop codons in two overlapping genes (*YNL114C* and *RPC19*) on chunk M3 resulted in a respiratory growth defect on YPG medium (Figure 5A). Closer inspection of these two genes revealed that the reassigned TAA stop codon of the dubious ORF *YNL114C* likely interfered with transcription of the antisense overlapping *RPC19* ORF, possibly by altering transcription factor binding. Subsequent integration of the *RPC19* TAA stop codon while retaining the native *YNL114C* TAG codon did not result in any fitness defect, confirming that the *YNL114C* TAA codon had caused the initial defect. This final synXIV strain had wild-type fitness on YPD and YP-glycerol medium at both 30°C and 37°C (Figure 5B). It is important to note that because *YNL114C* is a dubious ORF, the retention of its native TAG stop codon does not affect the Yeast 2.0 project goal of TAG codon reassignment.

**Figure 5.**
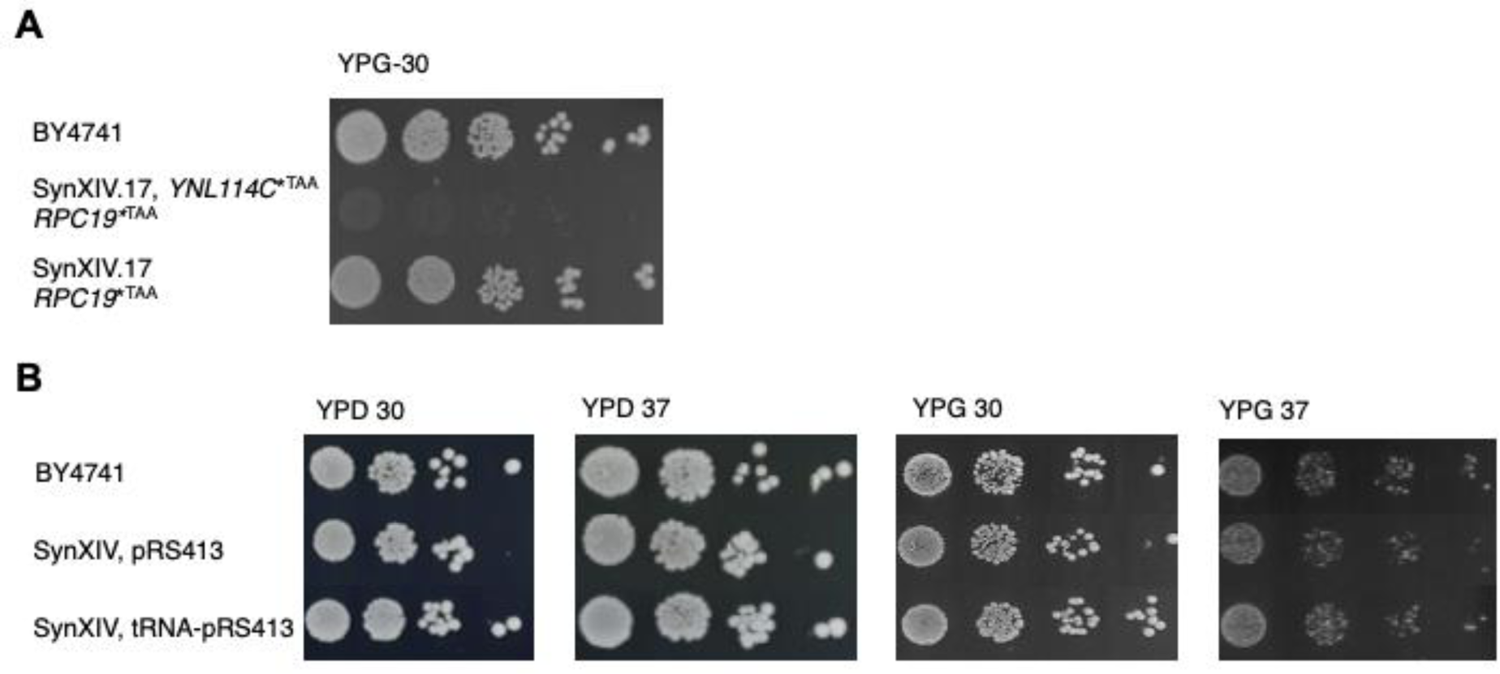
*YNL114C* dubious ORF stop-codon replacement causes a growth defect. (**A**) Replacement of the *YNL114C* TAG stop codon (strain 68, Table S1) but not the *RPC19* TAG stop codon (strain 69, Table S1) with TAA causes a growth defect on YP-glycerol medium relative to the wild-type control BY4741. (**B**) The fitness of fully synthetic chromosome XIV strains with and without deleted tRNA genes complemented on pRS413 (strains 77 and 76, Table S1) were compared with the wild-type strain (strain 75, Table S1). Images were taken after 4 d of growth at 30°C and are representative of two repeated experiments.

### Design and construction of synthetic-wild-type polyploid hybrids

The SCRaMbLE system is limited by the deletion of essential genes, and a subsequent reduction of viable cells in a population by over 100-fold after 24 h of induction (Shen *et al*., 2016). This limitation is particularly important when the phenotype of interest cannot be easily screened for, or when SCRaMbLE is used to incorporate foreign DNA, a process referred to as “SCRaMbLE-in” (Liu *et al*., 2018). In theory, SCRaMbLE-ing in heterologous DNA flanked by LoxP sites would enable integration of genes of interest in addition to synthetic genome rearrangements. Integration of heterologous genes is highly desirable for certain phenotypes such as cellulose degradation, where high concentrations of cellulase enzymes are required for optimal function (Kroukamp et al., 2017). We hypothesized that synthetic cells with higher ploidy would provide a viability buffer against the detrimental effects of essential gene loss, with the increased copy number of the synthetic chromosomes providing additional Cre-recombinase recognition sites for recombination, thereby enhancing the frequency of heterologous DNA SCRaMbLE-in efficiency. By sequential mating locus replacement and mating (Figure 6), strains with different combinations of native and synthetic chromosomes were isolated, except for a diploid and tetraploid strain exclusively harboring synthetic chromosomes III, VI and IX-R. This could be due to unintended changes to gene expression levels in diploid specific genes of the synthetic chromosomes (Strome et al., 2008).

**Figure 6.**
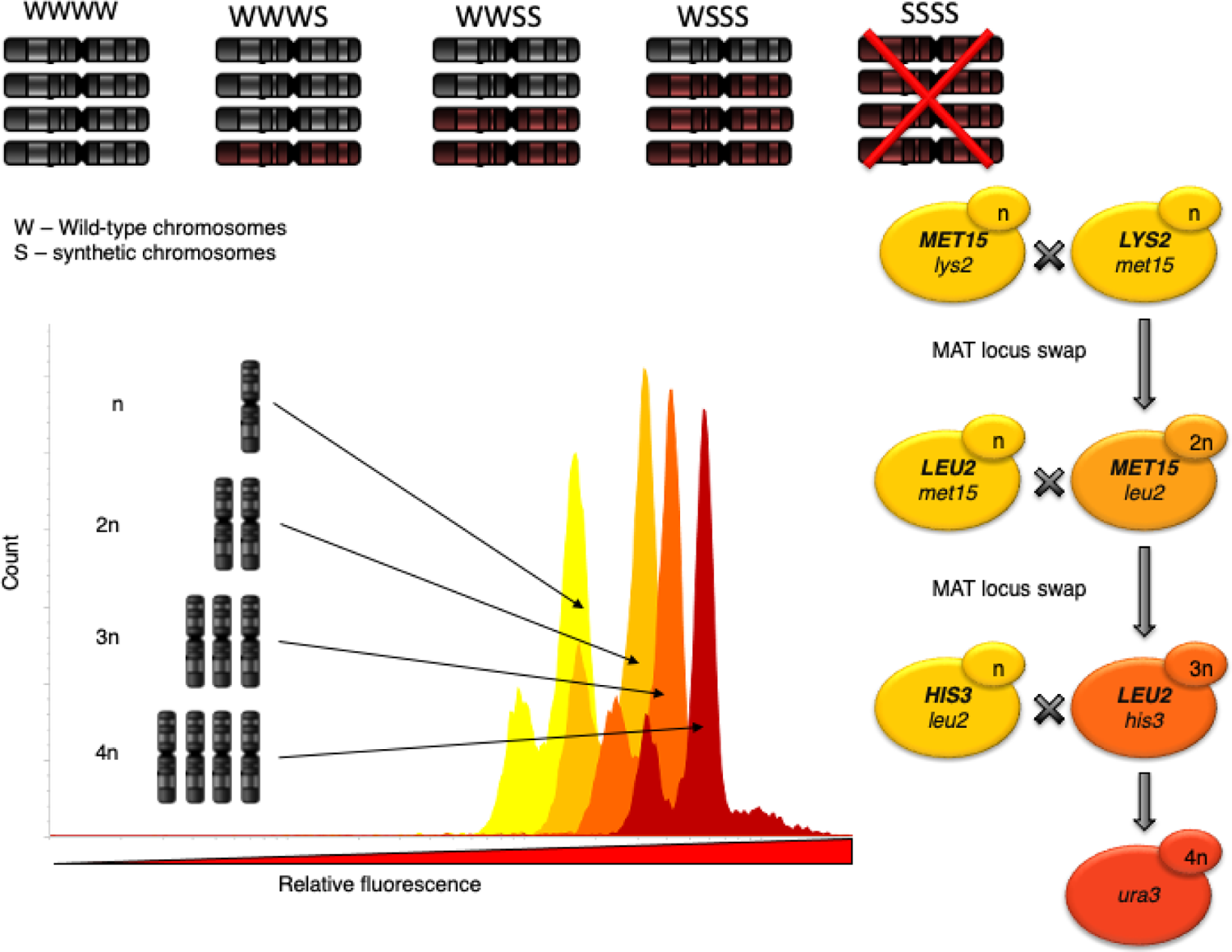
Polyploid strain construction strategy and ploidy verification. The leucine (*LEU2*) and methionine (*MET15*) auxotrophic markers were restored in *URA3* deficient *MAT***a** and *MAT*α versions of the wild-type and syn 3, 6, 9R strains (strains 79-84, Table S4), These were sequentially mated, resulting in haploid, diploid, triploid, and tetraploid strains with different combinations of wild-type (W) and synthetic (S) chromosomes III, VI, and IXR. Cells from each round of ploidy increase were selected on the appropriate nutrient deficient media based on the combined nutrient prototrophy created by the correct parent mating event. Diploid cells had their *LYS2* and *MET15* functionality restored, triploids had an additional *LEU2* restored, while tetraploid cells had an additional *HIS3* restored, resulting in cells with only ura3-auxotrophy. This was verified by PCR amplification of the *MAT* locus genes and all the relevant auxotrophic marker genes. To achieve mating of the polyploid strains, the strains mating type was made homozygous at each step by transforming with the required mating type replacement cassette. Ploidy was verified using propidium iodide DNA staining with reference to known haploid and diploid strains. We were unable to construct a fully synthetic tetraploid (red X).

SCRaMbLE-in of a LoxP-flanked *URA3* gene (Figure 7A) in haploid yeast cells with synthetic chromosomes III, VI, and IXR (Mitchell *et al*., 2017) did not result in a significantly greater amount of transformants relative to a non-SCRaMbLE control culture (Figure 7B), 2-sided student t-test with p > 0.05). However, when the same experiment was carried out using diploids that had synthetic and wild-type copies of chromosomes III, VI, and IXR, there was a dramatic increase in LoxP-*URA3*-LoxP integration relative to a no-SCRaMbLE control (Figure 7C). This indicated that the wild-type chromosome copies provided genetic redundancy to reduce the effect of essential gene loss, and that the creation of synthetic-wild-type hybrid polyploid strains may provide a mechanism for mitigating the limitations of haploid SCRaMbLE.

**Figure 7.**
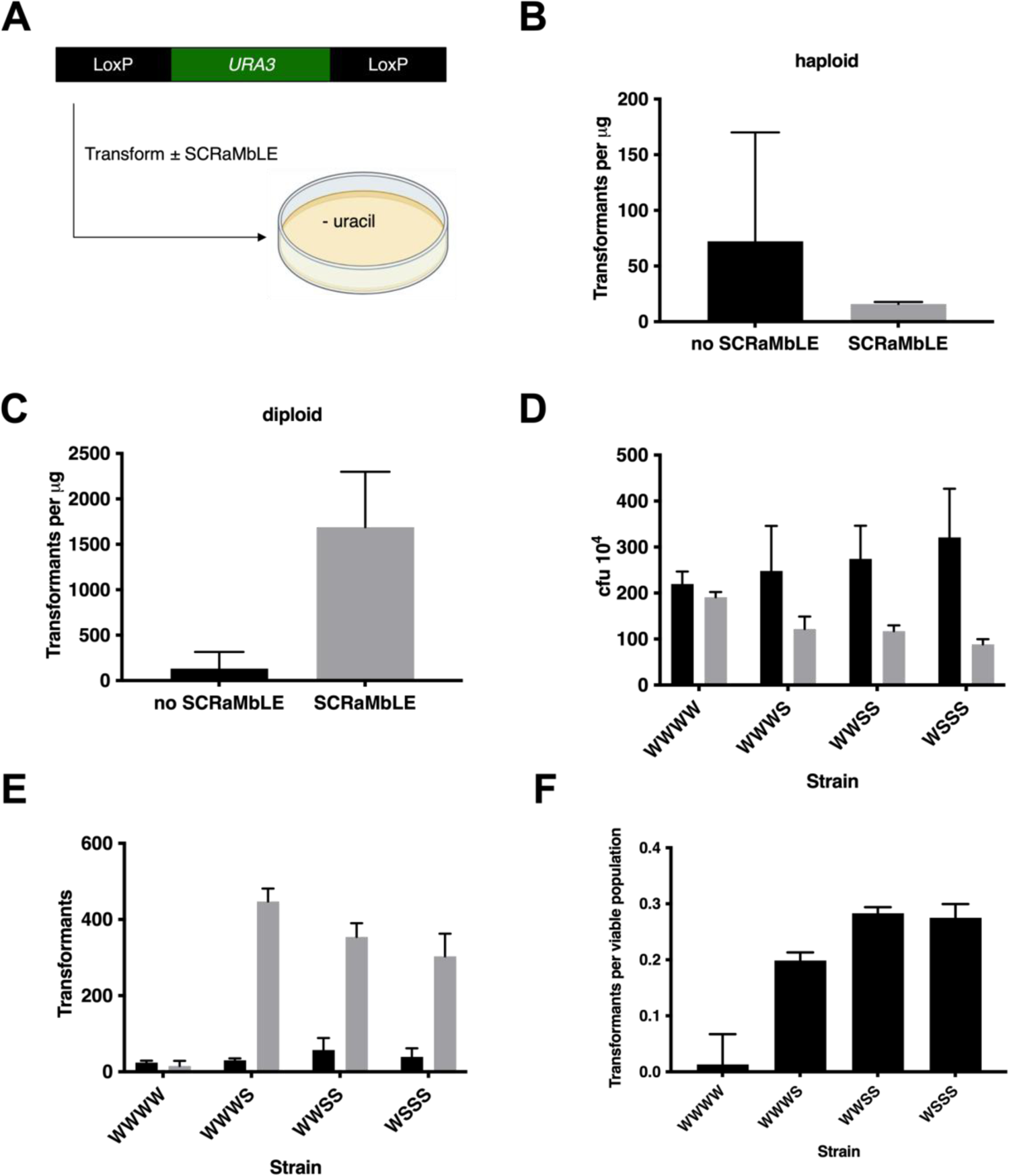
Variation of synthetic and wild-type chromosome ploidy for increased heterologous DNA SCRaMbLE-in efficiency. A loxPsym-flanked *URA3* expression cassette (A) was transformed into either haploid yeast with synthetic III, VI, and IXR chromosomes (Mitchell *et al*., 2017) (B, strain 80, Table S4), or a synthetic III, VI, IXR strain crossed with BY4741 to form a diploid (C, strain 86, Table S4). Tetraploid strains with different combination of synthetic (S) and wild-type (W) chromosomes II, VI, and IXR (strains 90, 91, 92, 93, Table S4) were used to measure viability with (gray bars) and without (black bars) SCRaMbLE-in from 90 µL of culture (D), LoxP-flanked *URA3* transformant number with (grey bars) and without (black bars) SCRaMbLE (E), and transformants per viable cell (F). Results from three independent transformations are shown with bars and error bars representing mean and standard deviation, respectively. SCRaMbLE-treated and non-SCRaMbLE populations were derived from the same transformed population. See also Figure S4.

To further explore the effect of synthetic-wild-type hybrid genome ploidy on SCRaMbLE, tetraploid strains (Figure 6) were tested for post-SCRaMbLE viability after 4 h of estradiol induction. Surprisingly, there was still a significant reduction in colony formation in synthetic chromosome-carrying strains relative to the wild-type control (Figure 7D, one-way ANOVA with Dunnet adjusted p-value for multiple comparisons to the fully wild-type tetraploid WWWW strain, p < 0.05), but the effect did not increase proportionally with the number of synthetic chromosomes present. This indicated there is still a significant viability loss that even in the presence of genetic redundancy in the form of wild-type chromosomes in polyploid strains. A possible explanation for this finding could be the generation of genetically unstable aneuploid strains or the loss of the *MATα* locus during the SCRaMbLE procedure, as we only rarely observed fully triploid strains during tetraploid strain generation.

In nature, diploid organisms are protected against essential gene loss by the presence of an extra copy of each gene, yet SCRaMbLE’d semi-synthetic diploid strains had a 40 % reduction in viability after 8 h of growth before recovery (Figure S4). The extra wild-type chromosomes increased the survival of the semi-synthetic diploid over a synthetic haploid strain which had a 60 % reduction in cell density after 10 h of Cre-recombinase induction. Although the S and WS strain population densities recovered over longer growth periods, the initial high cell death rates allow cells with limited SCRaMbLE events (which are more fit) to dominate, resulting in an undesired final population with low genotypic diversity (Wightman et al., 2020). The tetraploid strains displayed improved viability during SCRaMbLE, with the WSSS strain containing the largest number of synthetic chromosomes, only losing ∼30% viability.

## Discussion

We found that a respiratory growth defect was caused by the insertion of a loxPsym site 3’ of the mitochondrial ribosomal protein encoding *MRPL19* gene (Figure 1). Like all nuclear-encoded mitochondrial proteins, the *MRPL19* protein is targeted to the mitochondria. Consistent with this, the *MRPL19* protein sequence includes a predicted mitochondrial targeting peptide signal in the N-terminus (Fukasawa et al., 2015). In addition to protein targeting signals, many nuclear encoded mitochondrial genes have motifs in the 3′ UTR of their mRNA that facilitate localization to the outside of mitochondria via recognition by the *PUF3* protein for co-translation and import (Saint-Georges *et al*., 2008). It is noteworthy that the Puf3p binding motif (UCUGUAAAUA (Lapointe et al., 2018)) is located 81 bp 3′ of the *MRPL19* loxPsym site, meaning that the observed defect may not be mediated by disruption of Puf3p binding to *MRPL19* mRNA. Alternatively, Crg1p, Mtq2p, and Scd6p have been shown to bind *MRPL19* mRNA (Tsvetanova et al., 2010), and it is possible that they have a role in mediating the Mrpl19p mis-localization phenotype.

Backcrossing of the fully synthetic J1.8 strain to a wild-type followed by whole genome sequencing and fitness testing of haploid progeny led to the discovery of a series of background mutations on other chromosomes that impacted fitness (Figure 3). In particular, mutations detected in ATP synthase subunit genes were previously observed to suppress defects associated with mitochondrial DNA loss by allowing mitochondrial protein import in petite strains, yet would be deleterious when mitochondria are present and functional (van Leeuwen et al., 2019). The history of the synXIV strain involved a long period without mitochondrial DNA, during which time the observed *ATP1^A424T^* and *ATP3^I303V^* mutations may have arisen. When the two half-chromosome strains were crossed (Figure S1), we generated a strain with healthy mitochondria which lacked the synthetic DNA region that initially caused mitochondrial dysfunction (*MRPL19*) (Supplementary Figure 1C, Figure 2), meaning that the background *ATP1* and *ATP3* mutations may have switched from suppressing a growth defect to causing one.

Integration of missing synthetic DNA regions into the backcrossed synXIV.17 strain showed that deletion of the *NOG2* intron on megachunk W also resulted in a growth defect (Figure 4, Figure S2). Although *NOG2-GFP* fluorescence levels were similar with and without the *NOG2* intron in the SynXIV.17.W.X strain (Figure S2 C), growth was only restored to wild-type levels when the intron-containing wild-type *NOG2* gene was expressed from a plasmid (Figure S2 D). *NOG2* is an essential, intron-containing gene encoding a putative GTPase that facilitates pre-60S ribosomal subunit maturation and export from the nucleus (Saveanu et al., 2001). The *NOG2* intron encodes a small nucleolar RNA (snoRNA), which guides pseudouridylation of large subunit (LSU) rRNA (Badis *et al*., 2003) at positions that are highly conserved across bacterial and eukaryotic domains. These rRNA pseudouridylations facilitate the formation of correct ribosome structure, and the *snr191* snoRNA has previously been reported to convey a growth advantage in yeast, but is not essential (Badis *et al*., 2003). The linear *NOG2* intron also accumulates under stress conditions as a ‘stable’ intron, where it is disadvantageous if deleted (Morgan *et al*., 2019). It would be interesting to try to produce a “refactored” linear version of the snr191 snoRNA which would in principle allow the intron to be deleted without loss of fitness.

In parallel to our rational debugging approaches (Figure 3), the defective J1.8 strain had wild-type growth restored via ALE on YP-glycerol medium, which led to increased *TAR1* copy number (Figure 2). Nanopore sequencing showed that this increased *TAR1* copy number likely occurred in the form of ERCs, which are known to accumulate in yeast with defective mitochondria (Borghouts *et al*., 2004). *TAR1* expression at the rDNA locus is repressed by the *Sir* complex (Li *et al*., 2006), meaning that relocation of *TAR1* to ERCs could increase Tar1p expression to mediate mitochondrial activity (Poole *et al*., 2012). Recent work has shown that when defective ‘selfish’ mitochondria are inherited after mating, the retrograde response is triggered, leading to *TAR1* amplification to repress and remove defective mitochondria from the population (Poole *et al*., 2012; Walker, 2015). The increased rDNA copy-number we observed in evolved populations could have occurred in response to defective mitochondria that were inherited from the crossing of partially synthetic strains to make synXIV (Figure S1A). Defective mitochondria would have been present in the synthetic G-X strain for many generations after the introduction of megachunk J and the growth defect associated with the *MRPL19* loxPsym motif.

In retrospect, given the number of potentially deleterious background mutations (Figure 3) and synXIV design changes (Figures 4, 5) that had to be reverted to restore normal growth rationally, it makes sense that our parallel ALE approach resulted in a broad suppressor mutation in the form of *TAR1*-carrying ERCs. The increasing scale and complexity of synthetic genome projects means that synthetic lethality and synthetic growth defects will likely become more common, which will increase the difficulty of rational debugging approaches. ALE is therefore set to ultimately become an even more important tool in this field, where synthetic genomes will need to be debugged with limited prior knowledge of the genetic basis of bugs. ALE can also be used to ‘polish’ synthetic genomes towards improved industrial fitness, as recently demonstrated in a recoded *Escherichia coli* genome (Ostrov et al., 2016; Wannier et al., 2018). However, our results show that when multiple bugs are present, an ALE approach may result in broad suppressor mutations that are effective, yet do not fully resolve the underlying ‘original’ genetic basis of the poor growth phenotype. Whether evolved suppressor mutations can be tolerated will depend on individual project goals and parameters, and if rational debugging is technically feasible.

In addition to the construction and debugging of synXIV, we further developed the Sc 2.0 SCRaMbLE system to tolerate essential gene loss by engineering synthetic-wild-type tetraploid hybrid strains. The increased number of synthetic chromosomes within the tetraploid strains allowed a greater variety of recombination events through more LoxP sites to facilitate SCRaMbLE-in (Figure 7, Figure S3), while promoting the recovery of a more diverse post-SCRaMbLE cell population. These synthetic-wild-type tetraploid strains further enhance the genetic redundancy and therefore the potential genotypic diversity of SCRaMbLE’d populations, in line with previous work on synthetic-wild-type diploid strains (Jia *et al*., 2018; Liu *et al*., 2018). Together with our exploration of parallel rational and non-rational growth-defect debugging, our results on tetraploid SCRaMbLE-ing demonstrate the extreme plasticity of synthetic genomes to both designer and random changes in gene content and genomic architecture.

## Acknowledgements

The Synthetic Biology initiative at Macquarie University is financially supported by an internal grant from the University, and external grants from Bioplatforms Australia, the New South Wales (NSW) Chief Scientist and Engineer, and the NSW Government’s Department of Primary Industries. Ian Paulsen is supported by an Australian Research Council Laureate Fellowship. TCW and BL were supported by Fellowships from the CSIRO Synthetic Biology Future Science Platform and Macquarie University. TCW and ISP acknowledge the support of ARC Discovery Project DP200100717. BL acknowledges the support of the Gordon and Betty Moore Foundation (GBMF9319, grant DOI:https://doi.org/10.37807/GBMF9319). Work in the JDB lab was supported by US NSF grants MCB-766 1026068, MCB-1443299, MCB-1616111 and MCB-1921641. Work in the JSB lab was supported by US NSF awards MCB-1445545 and EF-1935355.

## Author Contributions

Conceived and coordinated the project (JDB, ISP, DLJ, YC, NC, TCW, ITP, TRM, LAM, JSB). Designed experiments (TCW, ITP, XX, HK, TRM, IWD, ARB, HKN, BL, NVW, ITP, RSKW, YC, HDG, GS, JSB). Conducted experiments (TCW, HK, XX, ELIW, MIE, IWD, ARB, BL, NVW, ELD, RSKW, ACC, HDG). Analyzed data (TCW, HK, XX, NVW, ARB, BL, ELIW, MIE, HDG, GS). Wrote the manuscript (TCW, HK, ITP, ISP, JDB). All authors revised and approved the final manuscript.

## Methods

### RESOURCE AVAILABILITY

#### Lead contact

Further information and requests for resources and reagents should be directed to and will be fulfilled by the lead contact, Tom Williams.

#### Materials availability

All plasmids and yeast strains generated during this study are available on request.

#### Data and code availability

- Genome sequencing data has been deposited at NCBI under BioProject ID PRJNA841391 and will be made publicly available as of the date of publication. Accession numbers will be listed in the key resources table. Microscopy data reported in this paper will be shared by the lead contact upon request.
- This paper does not report original code.
- Any additional information required to reanalyze the data reported in this paper is available from the lead contact upon request.

### EXPERIMENTAL MODEL AND SUBJECT DETAILS

*S. cerevisiae* strains (Table S1 and Table S4) are all derivatives of BY4741 (MATa *his3Δ1 leu2Δ0 met15Δ0 ura3Δ0*), a haploid auxotrophic laboratory strain of mating type ‘a’. Yeast strains were grown in synthetic dropout (SD) media containing Yeast Nitrogen Base Without Amino Acids mix (Sigma-Aldrich Y0626) supplemented with 10 g/L glucose, and amino acids at 100 mg/L to complement auxotrophies as appropriate. Alternatively, yeast were grown in 10 g/L yeast extract, 20 g/L peptone, 20 g/L dextrose (YPD), or YP-glycerol (20 g/L glycerol in place of dextrose). For strains expression β-glucosidase, YP-cellobiose selective plates were prepared (20 g/L cellobiose in place of dextrose). *E. coli* DH5α strains were used to store and propagate plasmids (Table S3), and were grown in Lysogeny Broth medium with 50 mg/mL ampicillin.

Liquid growth of *E. coli* and *S. cerevisiae* strains was carried out in an Infors 25 mm orbital shaking incubator set to 30°C or 37°C and 200 rpm. Cells were cultured in either sterile 50 mL falcon tubes or 250 mL baffled shake-flasks where medium did not comprise more than 10 % of the total vessel volume.

### METHOD DETAILS

#### Growth Media

*S. cerevisiae* strains were grown in medium containing synthetic complete (SC) media containing 1x Yeast Nitrogen Base Without Amino Acids mix (Sigma-Aldrich Y0626) supplemented with 10 g/L glucose, and amino acids at 100 mg/L to complement auxotrophies as appropriate. Alternatively, yeast were grown in 10 g/L yeast extract, 20 g/L peptone, 20 g/L dextrose (YPD), or YP-glycerol (20 g/L glycerol in place of dextrose).

#### Growth conditions

Liquid growth was carried out in a 25 mm orbital shaking incubator (Infors Multitron Pro) set to 30°C and 200 rpm. Cells were cultured in either sterile 50 mL Falcon conical tubes or 250 mL baffled shake-flasks where medium did not comprise more than 10 % of the total vessel volume.

#### Chunk preparation

DNA chunks comprising ∼5-10 Kb of each megachunk were synthesized and sequence verified by Genscript (megachunks A-K and N-X), GeneArt (megachunks L, M), and GeneWiz (chunks E1, E2, E3, S4). Chunks were then either restriction digested using terminal, complementary sites incorporated in the design changes, or PCR amplified using primers that anneal to the 5’ or 3’ ends of each chunk with Phusion polymerase (New England Biolabs). Plasmid digested or PCR amplified chunks were excised from agarose gels or column purified, and quantified. Chunks were pooled together such that the relative amounts of each chunk were approximately halved so that chunk 1 > chunk 2 > chunk 3 > chunk 4, with the amount of chunk 4 being 400-800 ng. Restriction digested chunks were ligated over night at 16°C using T4 DNA ligase (New England Biolabs).

#### Yeast transformation and marker-loss screening

Cells were transformed using the lithium acetate/polyethylene glycol/ssDNA transformation method (Gietz and Schiestl, 2007). After 2-5 d of incubation on selective media at 30°C, colonies were replica-plated onto media selective for the marker gene used to integrate the previous megachunk, with those not able to grow used for further analysis.

#### DNA extraction and PCR-tag analysis

Genomic DNA was extracted using a lithium acetate-SDS solution for cell disruption followed by ethanol mediated DNA precipitation as previously described (Lõoke et al., 2011). Crude DNA extracts were transferred to a 384-well plate compatible with the Echo 550 acoustic liquid transfer system (Labcyte), as were primer pairs for each PCR-tag in synXIV (File S2) (15 μM). 4.75 μL aliquots of 1x SYBR green mastermix were added to each well of a 384-well qPCR plate using an epMotion liquid handling robot. 200 nL of crude gDNA and 50 nL of each primer-pair was transferred to each 384-well qPCR plate well using an Echo550 (Labcyte Inc.). The plate was centrifuged briefly to ensure transferred droplets were suspended in the SYBR-green mix. qPCR was carried out using a Lightcycler 480 with an initial 95°C denaturation of 3 minutes followed by 15 cycles of 30 s at 95°C, 30 s at 70°C with a decrease of 1°C each cycle, and extension at 72°C for 30 s. The same denaturation and extension condition were then used for a further 20 cycles, except with constant annealing at 55°C. SYBR-green fluorescence was measured at the end of each extension step. After cycling, a melt curve was generated by heating from 50°C to 95°C with fluorescence measurements every 5 s. For each megachunk, positive and negative controls were used that comprised of mixed synthetic chunk DNA or BY4741 DNA respectively. Any PCR-tags resulting in aberrant amplification were excluded from analysis of transformant DNA. Megachunk integration was accepted when all synthetic PCR-tags and no wild-type PCR-tags resulted in amplification.

#### CRISPR-Cas9 mediated genome modification

CRISPR-Cas9 mediated homologous recombination was carried out by using a previously reported strategy that utilizes a single episomal plasmid (pRS423) that contains both guide RNA and Cas9 expression cassettes (Williams *et al*., 2017). New 20 bp guide RNA sequences were encoded in 5’ extensions of primers that target the 3’ end of the *SNR52* promoter (reverse primers) and the 5’ end of the structural CRISPR RNA (forward primer). ∼ 100 ng of the linear PCR product resulting from this reaction was used to co-transform yeast with 1-5 μg of donor DNA with homology to the target guide-RNA locus. Colonies growing on SC –histidine media were screened for desired mutations using PCR-tag analysis and/or loci specific primers.

#### Fitness testing

Strains were inoculated into 5 mL of medium in 50 mL Falcon conical tubes and grown overnight at 30°C with 200 rpm shaking. Each culture was then passaged to a fresh tube with 5 mL of medium at an A_600nm_ of 0.4 – 0.5 and grown for a further 3-4 h. The A_600nm_ of each culture was adjusted to be the same, and each culture was 10-fold serially diluted in sterile MilliQ water down to 10,000-fold. 3 uL of each dilution was then spotted onto the indicated agar plates and incubated at 30 or 37°C for 4 d. Plates were imaged using a Singer Phenobooth, contrasts adjusted in Microsoft Powerpoint, and each dilution series cropped, resized, and repositioned without any non-proportional resizing. Only cultures that were grown on the same plate for the same amount of time were directly compared and shown together. Each image is representative of at least two repeated experiments. Strains harboring plasmids were precultured in appropriate selective liquid media.

#### SynXIV discrepancy repair

As a default option, sequence discrepancies (Table S2) were repaired using our previously developed CRISPR-Cas9 system (Williams *et al*., 2017), whereby synthetic chunk DNA served as donor for homologous recombination. Discrepancies 1-10 (Table S2) were repaired by targeting the *EGT2* ORF with CRISPR-Cas9 and synthetic chunks A3-A4 as donor DNA. The *EGT2* ORF was fully synonymously recoded as part of an error in the synXIV design phase, and we subsequently reverted this sequence to wild-type. However, this made no difference to fitness, so the wild-type sequence was used as a guide-RNA target during the repair of discrepancies 1-10, leaving the sequence in its original fully recoded state. Two non-TAA stop codons on the *YNL114C* and *RPC19* genes on chunk M3 were re-inserted by integrating a chemically synthesized *URA3* marker flanked by 796 bp 5′ and 1236 bp 3′ homology to the region. The *URA3* marker was then replaced by homologous integration of donor DNA with either both stop codons swapped, or with only the *RPC19* stop codon swapped (discrepancy 17, along with 18) using a *URA3*-specific CRISPR guide RNA and Cas9. Discrepancies 15, 16, and 19 were repaired using a similar approach, whereby a synthetic *URA3* marker was inserted, then replaced with PCR amplified DNA containing the desired changes. Discrepancies that were located on terminal, marker-containing or overwriting chunks (numbers 11-14 and 20-23, Table S2) were repaired by integrating the relevant chunk using selection for its marker (*LEU2* or *URA3*). The marker was then removed by targeting it using a *LEU2* or *URA3* specific CRISPR-Cas9 cassette and 3′ chunk donor DNA. Primers used to amplify the CRISPR-Cas9 cassette and encode *URA3* or *LEU2* specific guide RNAs, with guide RNA sequences in lower case (URA3 crRNA F: agcttggcagcaacaggactGTTTTAGAGCTAGAAATAGCAAGTTA URA3 crRNA R: agtcctgttgctgccaagctGATCATTTATCTTTCACTGCG, LEU2 crRNA F: ggcaacaaacccaaggaaccGTTTTAGAGCTAGAAATAGCAAGTTAAA, LEU2 crRNA R: acggttccttgggtttgttgccGATCATTTATCTTTCACTGCGGA). A duplication between the *ECM22* and *HAP1* genes on chromosome XII was discovered after sequencing of strain 55. This discrepancy was repaired by inserting a *URA3* marker cassette immediately after the stop codon of *EMC22*, growing without *URA3* selection overnight in YPD medium, plating for single colonies, and replica plating onto YNB glucose medium with 5FOA to select for colonies that had randomly looped out the chromosome XII duplication. Colonies were screened for loss of the *URA3* marker using PCR, and for single copies of the *SYM1* and *EST1* genes using RT-qPCR on genomic DNA with the *ALG9* gene used for normalization as previously described (Williams *et al*., 2015). Removal of this duplication was subsequently confirmed using whole-genome sequencing, and had no effect on strain fitness. Fully synthetic versions of synXIV that were whole genome sequenced are described in Table S5.

#### *MRPL19*-GFP fusion protein design

File S4 contains annotated genbank files of the plasmids and genes for the *MRPL19* protein internally tagged with super-folder Green Fluorescent Protein (sfGFP) (Pédelacq et al., 2005) into its coding sequence with and without the loxPsym site in the 3′ UTR, and a cytosol-localized GFP control (Williams *et al*., 2017), respectively. We inserted an in-frame *sfGFP* sequence inside the coding sequence of *MRPL19* (between position 282 and 283) because this gene encodes a predicted mitochondrial N-terminal peptide targeting signal (Fukasawa *et al*., 2015) and a 3′ UTR mRNA signal that mediates mRNA localization to mitochondria-bound polysomes involved in mitochondria protein import (Saint-Georges *et al*., 2008). The mitochondrial N-terminal peptide targeting signal was identified using MitoFates (Fukasawa *et al*., 2015) and by generating and analyzing a 3D protein model of Mrpl19p using SWISS-MODEL (Waterhouse et al., 2018), which also revealed an N-terminal β-hairpin motif predicted to target proteins to mitochondria (Jores et al., 2016). These mitochondrial targeting signals would have been disrupted by placing the sfGFP at either the N- or C-terminus of Mrpl19p. To promote proper folding of this fusion protein, we flanked the sfGFP with flexible linkers (L) (Edwards et al., 2008) halfway through the *MRPL19* ORF. The resulting fusion protein had the following design: *MRPL1994-L-sfGFP-L-95MRPL19*. The native promoter and terminator regions were maintained, except for the version containing a loxPsym site 3 bp after the stop codon. These two cassettes were synthesized by Genscript Inc. and cloned onto pRS416 vectors using XhoI and NotI.

#### *TAR1* expression construct cloning

The *TAR1* gene and its native promoter and terminator were synthesized as an IDT gBlock and cloned onto the pRS413 vector using BamHI and NotI restriction sites. The annotated vector map is included as File S5.

#### *NOG2-GFP* expression construct design and cloning

Expression constructs for *NOG2-GFP* fusion genes were synthesized by Genewiz and cloned onto pRS416 using. The native *NOG2* promoter and terminator were used, and two versions were made, with and without the *snr191* encoding *NOG2* intron sequence. Annotated genbank files of these two plasmids are included in File S6.

#### tRNA-array design and cloning

As per Sc2.0 design principles, all tRNA genes are to be relocated (Richardson *et al*., 2017). To complement their loss from SynXIV, the synthetic ∼9kb ChrXIV tRNA array was designed to house all 14 tRNA genes relocated from the wild-type chromosome XVI of *S. cerevisiae*. Each tRNA gene was assigned 500 bp 5’ and 40 bp 3’ flanking sequences recovered from the yeasts *Ashbya gossypii* or *Eremothecium cymbalariae* to reduce homology to the host genome. tRNA flanking sequence assignment was automated using Python programming scripts based on an algorithm that matched tRNA genes to their flanking sequences preferentially by anticodon, and additionally altered unwanted artefacts (such as transcriptional gene starts) from the 5’ flanking sequence. Furthermore, rox recombination sites were designed to be placed at 5’ and 3’ intervals and all tRNA introns were removed. Following synthesis (Wuxi Qinglan Biotech Co. Ltd (Yixing City, China)), the ChrXIV tRNA array was clone into a pRS413 centromeric vector with NotI restriction sites introduced for subsequent removal (File S7). There are no single-copy or otherwise essential tRNAs in this array.

#### Confocal microscopy

BY4741 strains transformed with *MRPL19-sfGFP-loxP-pRS416*, *MRPL19-sfGFP-Native 3′ UTR-pRS416,* or cytosol localized GFP expression (*pPDR12-GFP-pRS416* (Williams *et al*., 2017)) were pre-cultured twice in minimal medium without uracil before being inoculated at an A_600nm_ of 0.4 in fresh medium. Cells were treated with 100 nM Mitotracker Red FM (ThermoFisher M22425) for 3-4 h with shaking at 30°C. Cells were kept on ice prior to microscopic examination. Visualization of GFP and Mitotracker Red FM signals was performed using an Olympus FV 1000 confocal laser-scanning microscope. Microscopy images were analyzed using ImageJ (https://imagej.nih.gov/ij/index.html). Images shown are representative of cells in independent biological triplicate populations.

#### Diploid formation

Strains of opposite mating type and with complementary auxotrophies were grown overnight separately in 5 mL of selective SD media. 500 μL of each culture was used to inoculate the same non-selective 5 mL of SC medium, which was incubated overnight at 30°C without shaking. The overnight culture was washed twice in sterile MilliQ water before being plated on solid medium selective for the respective auxotrophies in each strain, such that only diploids would form colonies. Putative diploid colonies were checked using ‘mating type’ primers (Key Resources Table) to verify the presence of both ‘a’ and ‘alpha’ alleles at the *MAT* locus, indicating the formation of a diploid.

#### Sporulation, random spore isolation, and random spore screening

To initiate sporulation, diploid colonies were grown overnight in 5 mL of selective liquid SD medium, washed once with sterile MilliQ water, and plated on 10 g/L potassium acetate medium. Plates were incubated in the dark at room temperature for 4-7 d. Once asci were visible under light microscopy, as many cells as possible were scraped from the potassium acetate plate and resuspended in 500 μL of sterile MilliQ water with 10 units of zymolyase and 20 μL of beta-mercaptoethanol. This solution was incubated at 37°C for 3-4 h before being transferred to a 250 mL flask containing 20 mL of 425-600 μm glass beads (Merck G9268) and 30 mL of sterile MilliQ water. Flasks were incubated at 30°C overnight with 200 rpm shaking. The liquid fraction was recovered, washed once in sterile MilliQ, and a dilution series down to 10^-3^ plated on YPD with incubation at 30°C for 1-2 d. Colonies were replica plated onto SD plates selective for each of the auxotrophic markers present in the haploid parent strains, and any colonies found not growing on each plate type were selected for PCR-tag analysis using the 2^nd^ tag of each of the 22 synXIV megachunks. Colonies with synthetic PCR-tag amplification and without wild-type PCR-tag amplification were deemed likely to contain the corresponding megachunk, and further screened using all PCR-tags for the relevant megachunks.

#### RNA extraction

1.5 mL samples of mid-exponential phase cultures (A_600_ of 0.5 – 2.5) were pelleted by spinning at 12,000 x g for 2 min and removing the supernatant. Pellets were resuspended in 1 mL of RNAlater (ThermoFisher Scientific catalog number AM7020) and stored at −20°C. RNA was extracted after pelleting cells and removing RNAlater solution using the Zymo Research YeaStar RNA extraction kit (catalog number R1002) according to the user manual. Co-purified DNA was removed from RNA extracts using TURBO™ DNase (ThermoFisher Scientific catalog number AM2238) according to the user manual.

#### RT-qPCR

100 – 1000 ng of purified RNA was used for reverse transcription using an 18 nucleotide poly-T primer and SuperScript™ III Reverse Transcriptase (ThermoFisher Scientific 18080093) according to the user manual. A no-enzyme control was included for each RNA sample and subsequently used for qPCR to verify that no genomic DNA was contributing to cDNA concentration estimates. Reverse transcribed samples were diluted 1:100 in MilliQ water prior to qPCR analysis. Relative expression was performed using the modified-Livak method (amplification efficiency measured for each primer-pair and not assumed to be log2) with *ALG9* as a housekeeping gene (Teste et al., 2009), as previously described (Williams *et al*., 2015).

#### Whole-genome sequencing

A yeast genomic DNA extraction kit (ThermoFisher catalog number 78870) was used to isolate DNA according to the manufacturer’s instructions. Sequencing and library preparation were carried out by Macrogen Inc. using a True-Seq Nano kit with 470 bp inserts, and paired-end Illumina HiSeq 2500 sequencing, or by the Ramaciotti Centre for Genomics using Nextera XT library preparation and 2x 150 bp paired end sequencing using a NextSeq500 (Sequencing of samples from the adaptive laboratory evolution experiment). Reads were analyzed using Geneious Pro v10.2.2 software (Kearse et al., 2012). Paired-end reads were mapped to an edited version of the S288C reference genome where native chromosome XIV was replaced with synthetic chromosome XIV (File S7). The Geneious alignment algorithm was used to map reads to the reference genome using default settings. Analysis of the resultant assembly was completed visually by assessing read coverage, and read disagreement with the reference sequence. The raw reads were of high-quality (Q30 > 91 %, Q20 > 95 %), and were therefore not trimmed prior to assembly. Average read depth of 190 was typically achieved from the Macrogen sequencing, while 50-fold coverage was used for the samples sequenced at the Ramaciotti Centre for Genomics. Single Nucleotide Polymorphisms (SNPs) and their effect on ORF reading frames and codons were detected using the Geneious “Find Variations/SNPs” function with a variant p-value threshold of 10^-6^ and variant frequency of ≥ 50 %.

#### Nanopore sequencing

YP-glycerol evolved lineages had genomic DNA extracted as for Illumina sequencing. Barcoded nanopore sequencing libraries (SQK-LSK109) were prepared according to the manufacturer’s instructions and sequenced on a single flowcell (FLO-MIN106) using a MinION sequencer. Basecalling of raw FAST5 files was performed using albacore (v2.3.1), with subsequent barcode demultiplexing using Porechop (v0.2.3). Demultiplexed reads in fastq format were assembled with Canu (v1.7.1), with initial assemblies for each strain polished using nanopolish (0.10.1). Assembled contigs were annotated using the Geneious Prime ‘annotate from file’ feature, with the S288C genome with SynXIV used as a reference.

The penultimate synXIV strain (74, Table S1) was whole-genome sequenced after DNA extraction as above. Genomic DNA (1-2 µg) was prepared for ligation sequencing (SQK-LSK109) with native barcoding (EXPNBD104 and EXPNBD114) as per the manufacturer instructions. Following preparation, 200-300 ng of DNA was loaded onto a MinION flowcell (FLO-MIN106) and basecalling was performed with Guppy v4.0.11 or v4.2.3 (Oxford Nanopore Technologies).

#### Flow cytometry

GFP was measured using exponentially growing cultures at an A_600_ of 0.5 using a Becton Dickinson Accuri C6 flow cytometer. GFP fluorescence was measured using a 488 nm laser and a 533/30 emission filter. Mean GFP values were divided by the mean autofluorescence of an empty vector control strain.

#### Polyploid strain construction

Polyploid strains were constructed through sequential rounds of synthetic mating type switching and strain mating. In short, *his3* and *leu2* auxotrophies were complemented in independent synthetic yLM896 and BY4742 strains. This was achieved by PCR amplification of the relevant gene loci from extracted prototrophic S288c genomic DNA using the sets his3up-F/R and leu2up-F/R primers (Key Resources Table), and subsequent transformation into the respective strains to effectively restore the function of each respective auxotrophic loci. The relevant genotypes of the auxotrophic complementation strains are given in Table S4.

The WS and WW diploid strains were generated by co-inoculation of 1 mL of an exponential BY4741(k) culture (diluted to an A_600_ of 0.5) and 1 mL of an exponentially grown yLM896 or BY4742 strain, into 5 mL of YPD media and grown for 16 h at 30°C. The cultures were streaked out on SC –Met medium supplemented with 200 µg Geneticin to isolate single colonies. Diploid colonies were confirmed by the presence of both mating type loci via PCR amplification using the *MAT**a***, *MAT*α and MATlocus primers (Key Resources Table).

Diploid strains with heterozygous *MAT* loci are unable to mate spontaneously. To facilitate intermediate triploid strain generation, chemically synthesized *MAT**a*** DNA was transformed into the WS and WW strains. This allowed the generation of a small number of homozygous *MAT**a*** diploids within the population. After the heat shock step, transformed WS cells were co-inoculated with BY4742(L) cells and statically grown overnight in 5 mL fresh YPD medium at ambient temperature. The cell pellet was washed with sterile water, and the cell suspension was diluted and spread out to isolate single colonies on SC –Met – Leu agar plates. Putative triploid colonies resulting from mating were confirmed by the presence of both mating type loci via PCR amplification and named WWS. The same procedure was followed to generate the WWW and WSS strains, by mating the WW x BY4742(L) and WS x yLM896(L) strains respectively. The triploid genomic nature of each strain was verified through propidium iodide staining of nucleic acids and flow cytometry analysis.

The WWW, WWS, and WSS strains displaying a triploid genomic profile were selected for a subsequent round of strain transformation and mating as described above. The triploid strains had a *MATα/MAT**a**/MAT**a*** active mating locus genotype, allowing the conversion of a small number of cells within the population to homozygous *MAT**a*** strains. After transformation, cell pellets were washed and combined with either yLM896(H) or BY4742(H) to allow mating during the stationary overnight incubation. Putative tetraploid colonies were selected on SC -His -Leu agar plates. The tetraploid strains resulting from mating between WWW x BY4742(H), WWS x BY4742(H), WWS x yLM896(H) and WSS x yLM896(H) were selected based on their DNA content flow cytometry profiles and ability to grow on selective plates without leucine, histidine, methionine and with 200 µg/mL Geneticin (data not shown). The verified strains were designated WWWW, WWWS, WWSS and WSSS respectively.

For subsequent SCRaMbLE and growth characterization, the four tetraploid strains, and the haploid (W, S) and diploid (WW, WS) strains were transformed with pHK-Cre-EBDh, containing the Cre-EBD fusion-protein expression cassette. Strains containing the pHK-Cre-EBDh were selected on YDP agar plates supplemented with 200 µg/mL hygromycin B (Invivogen, USA).

#### Growth analysis of the tetraploid strains

To evaluate the effect of increased synthetic chromosomes on cell growth, the optical densities of W, S, WW, WS and the four tetraploid strains (containing the pHK-Cre-EBDh plasmid) were measured over time. Baffled flasks containing 15 mL fresh YPD containing 200 µg/mL hygromycin B were inoculated in triplicate to an A_600_ of 0.2 using stationary cultures of each respective strain. These cultures were incubated at 30°C with shaking at 250 rpm (Infors Multitron Pro). Optical density samples were taken every 2 h, after the first 8 h of growth.

The viability of each strain after inducing SCRaMbLE through estradiol activated Cre-recombinase expression was also evaluated. In parallel to the growth analysis, W, S, WW, WS and the four tetraploid strains were grown as described above, except with the addition of estradiol (Sigma-Aldrich) at t = 0 to achieve a final concentration of 1 µM. Viability was reported as the difference between the estradiol untreated and treated samples of the same strain at each given time point.

#### SCRaMbLE-in

Gene cassettes for the expression of *URA3* were synthesized with loxP sites flanking the cassettes. 1 µg of *URA3* cassette was transformed using the LiAc/PEG into the S and WS strains containing the Cre-expression vector, pLM006. After heat shock, cells were washed in SC -His media, and resuspended in either 5 mL of SC -His media or SC-His medium supplemented with estradiol to achieve a final concentration of 1 µM. Cells were recovered in these media for 1 h, washed with sterile water and plated on SC -Ura media. After heat shock, cells were washed in YPD media, and resuspended in either 5 mL of YPD + hygromycin media or YPD + hygromycin media supplemented with estradiol to achieve a final concentration of 1 µM. Cells were recovered in these media for 1 h, washed with sterile water and plated on selective plates. All transformations were done in triplicate. Colony forming units were counted from plating 90 µL

#### Ploidy determination

The relative cell DNA content determination protocol was adapted from Rosebrock (Rosebrock, 2017). Overnight cultures were inoculated into fresh growth media to an A_600_ of 0.2 and were grown to mid-exponential phase. Adequate cell culture was harvested to obtain 500 µL of culture at 2×10^7^ cells/mL. The cell pellet was washed with ice-cold water, and then fixed in 500 µL of ice-cold 70 % EtOH and incubated for at least 16 h at −20°C. The pellet was resuspended in Tris/MgCl_2_-buffer (50 mM Tris-HCl, pH 7.7, and 15 mM MgCl_2_) supplemented with RNase A to achieve a final concentration of 1mg/mL and incubated at 37°C for 90 min with gentle shaking. The pellet was then resuspended in 100 µL of 0.05 mM propidium iodide (PI) in Tris/ MgCl_2_-buffer and allowed to strain for 48 h at 4°C. The sample was then diluted and analyzed with a BD FACSAria using the 488nm laser for PI excitation and the PE filter to measure red light emission. PI stained BY4742 and BY4743 were used as haploid and diploid control samples, respectively. The BY4742 G1 cell cycle fluorescence peak was used as reference to estimate a haploid DNA complement, and its G2 peak as estimate for a diploid DNA complement. The BY4743 G2 peak was used as estimate for cells with a tetraploid DNA complement. Cells with a G1 peak that corresponded with the BY4743 G2 peak were considered to have a tetraploid DNA content, while cells with a G1 peak intensity in between the G1 and G2 peaks of the BY4743 strain were considered triploid.

### QUANTIFICATION AND STATISTICAL ANALYSIS

Statistical analysis was performed using Prism 9 and Microsoft Excel software. All of the statistical details of experiments can be found in the figure legends and results, including the statistical tests used, exact value of n, what n represents, definition of center, and dispersion and precision measures. Significance was defined using p-values of less than 0.05 with the tests indicated in the results section, no data or subjects were excluded.

---

**Supplementary S1.**
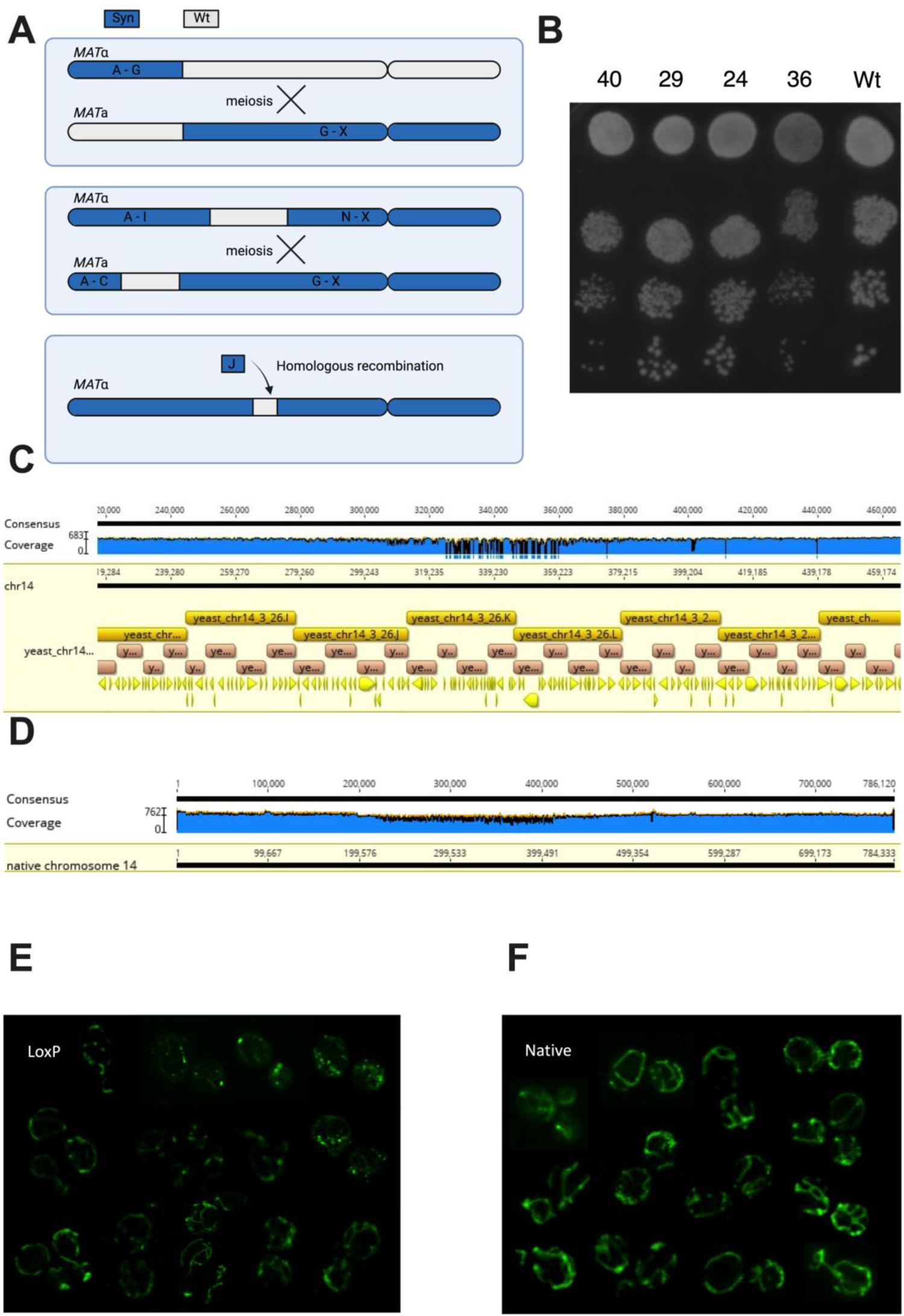
Pooled backcross sequencing and synthetic chromosome strain crossing reveals a fitness defect in megachunk J. (**A**) Megachunks A to X were integrated across two separate strains in parallel, with *URA3* and *MET17* markers integrated at the *YNL223W* and *YNR063W* wild-type loci of the synthetic A-G strain (strain 31, Table 3), respectively, prior to crossing with a synthetic G-X strain (strain 22, Table 3). Haploid strains resulting from the first cross (34 and 35, Table 3) were crossed together with screening for *LEU2* (*YNL125C* locus) and *URA3* marker (*YNL269W* locus) loss from wild-type regions to give a fully synthetic version of chromosome XIV. (**B**) Haploid progeny of a meiotic cross between two partially synthetic versions (A-G and G-X) of synXIV (colonies 40, 29, 24, 36) were tested for fitness alongside a BY4741 control (Wt) on YPD agar at 30°C for 3 d. Colonies 40 and 36 contain fully synthetic chromosome XIV, while colonies 29 and 24 contain synthetic DNA in all megachunk regions except I-J and J respectively. (**C**) Pooled sequencing of ‘fast’ growing haploid progeny of a cross between synthetic chromosome XIV strain G-X and BY4742. Each set of reads were aligned to a reference S288c genome that also had a copy of the synthetic chromosome XIV sequence. (**D**) ‘Slow’ reads mapping to native chromosome XIV. Dark yellow annotations correspond to megachunk regions while pink represent chunks, and yellow coding sequences. Average sequence coverage was 183 for the ‘Fast’ pool and 218 for the ‘Slow’. Chromosome annotation and coverage graphs were generated using Geneious Pro Software. (**E**) Versions of the *MRPL19*-GFP fusion genes were deigned with LoxP and (**F**) without LoxPsym motifs in the 3’ UTR GFP and expressed from pRS416 in the wild-type BY4741 strain. GFP fluorescence and distribution was visualised using an Olympus FV 1000 confocal laser-scanning microscope. Microscopy images were analysed using ImageJ (https://imagej.nih.gov/ij/index.html). Images shown are representative of cells in independent biological triplicate populations.

**Figure S2.**
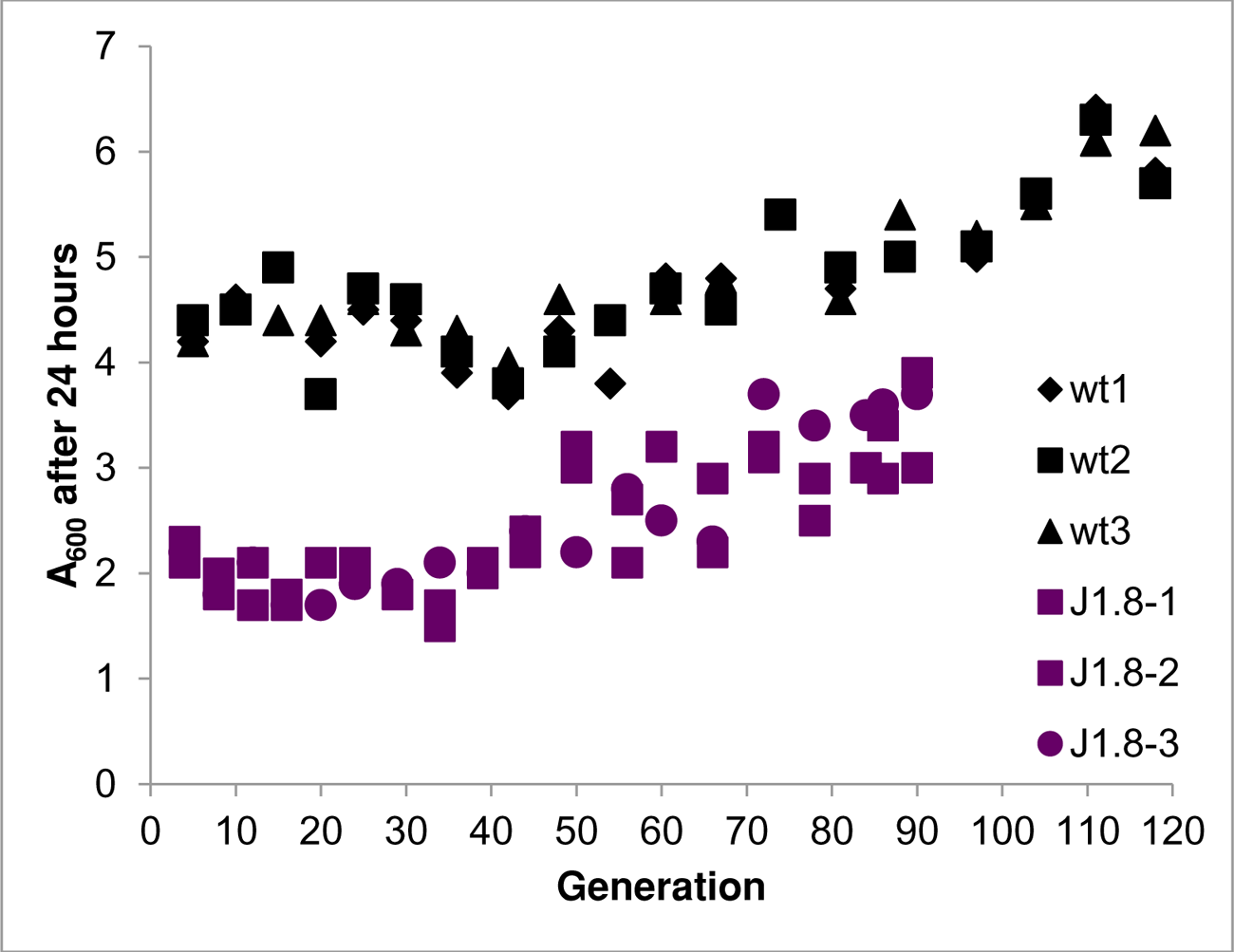
Fitness tracking of Adaptive Laboratory Evolution strains. A600 of three independent lineages for the BY4741 wild-type (strain 1 Table S1) and syn14 J1.8 strains (strain 40, Table S1) was measured every 24 h to track fitness and generation numbers in YP-glycerol medium.

**Figure S3.**
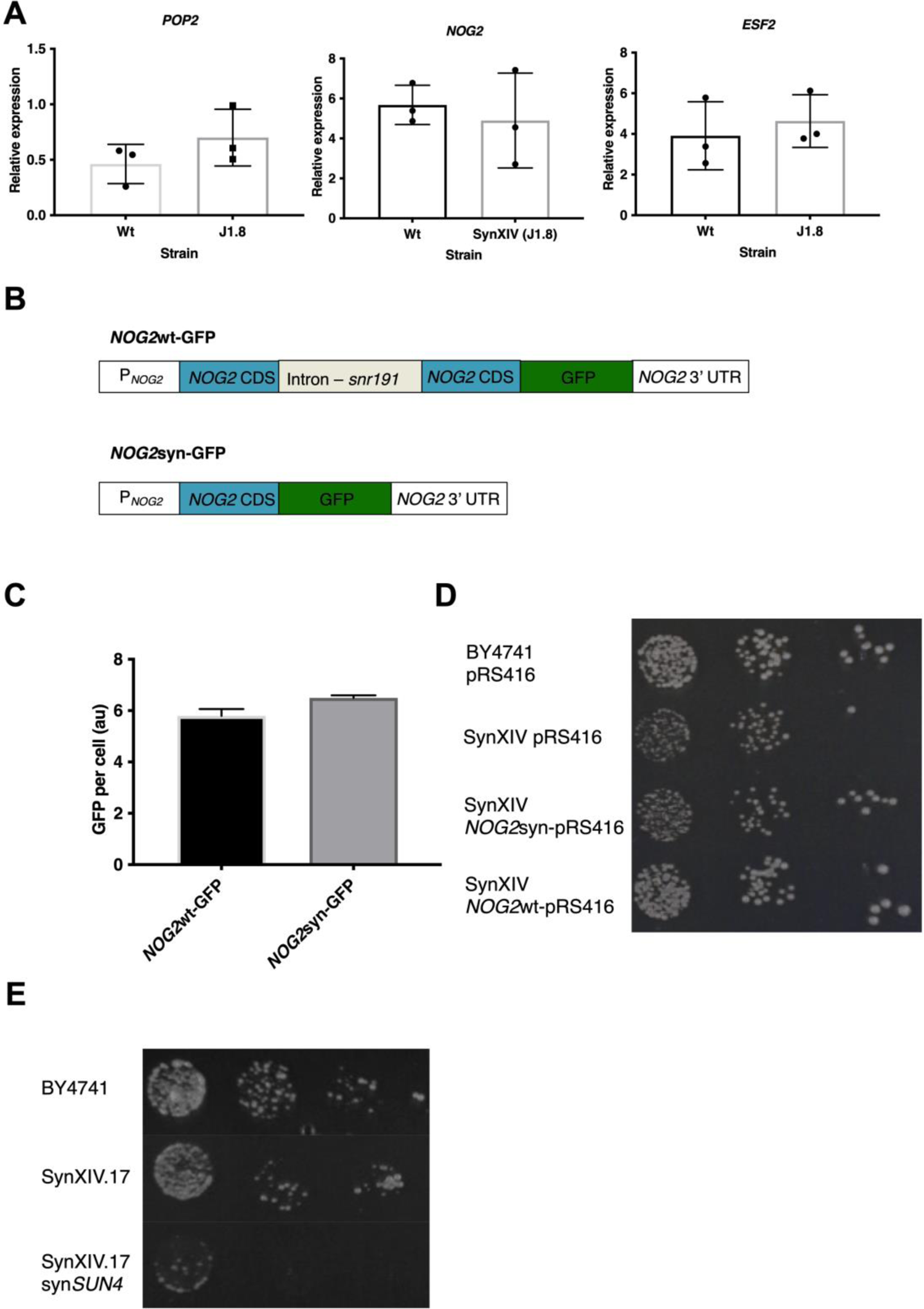
Expression of the *NOG2* and *SUN4* introns restores fitness. (**A**) mRNA levels of the three genes on chunk W1 were quantified using RT-qPCR. (**B**) Synthetic NOG2-GFP fusion expression constructs with and without the intron encoded *snr191* gene were designed and (**C**) tested for changes in protein expression the wild-type (strain 1, Table 3). (**D**) Expression of *NOG2-GFP* with its intron (strain 61, Table 3) but not without (strain 62, Table 3) restores growth to wild-type levels in synXIV on YP-glycerol medium at 37°C. Images were taken after three days. RT-qPCR and GFP fluorescence levels are reported as the mean of triplicate cultures with error bars plus or minus one standard deviation. (**E**) Serial 10-fold dilutions of wild-type (BY4741), SynXIV (strain 63, Table 3), and SynXIV with the *SUN4* intron removed (strain 64, Table 3). Strains were plated on YP-glycerol (YPG) at 37°C for 4 d prior to imaging.

**Figure S4.**
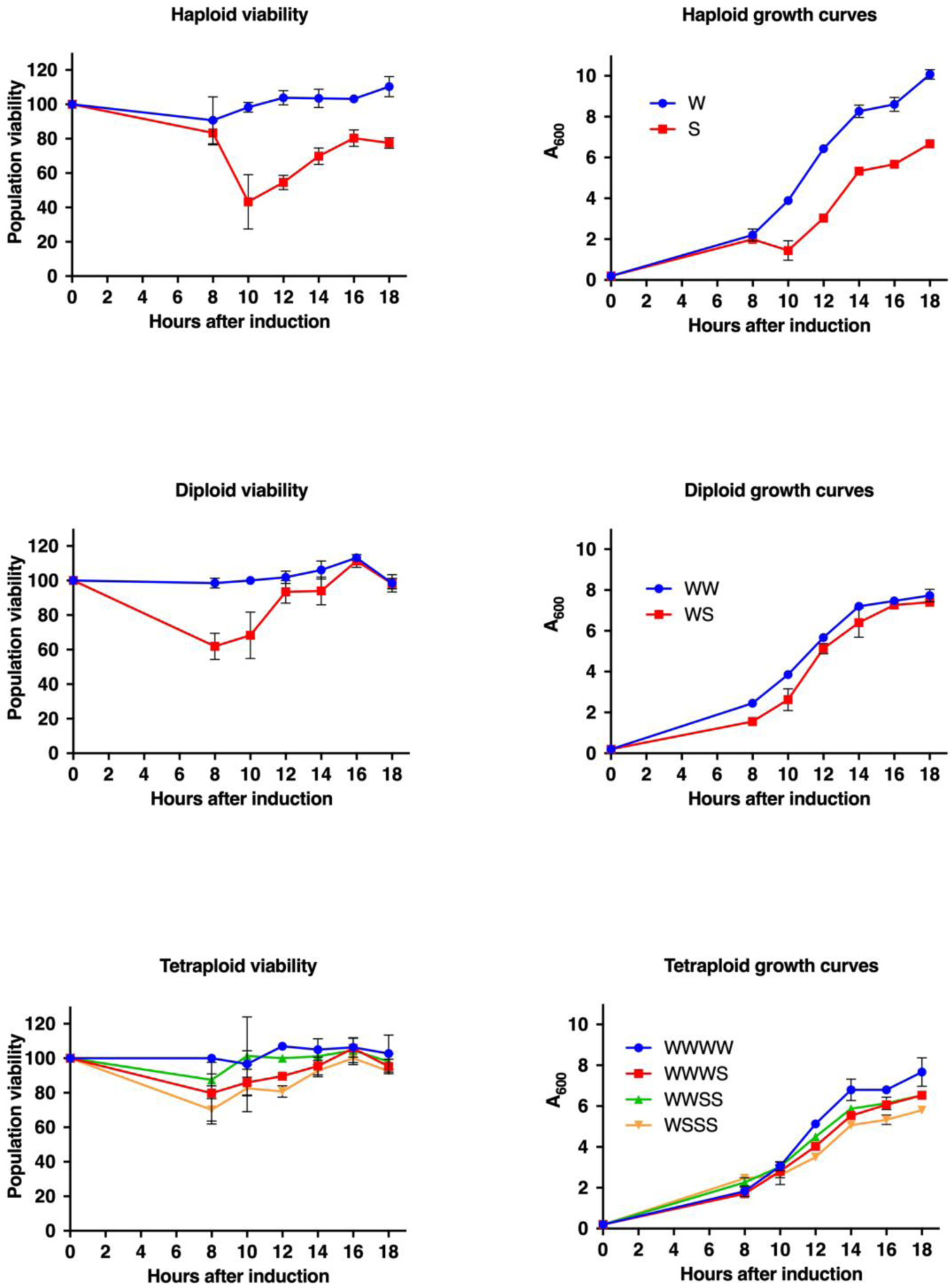
Growth and viability after SCRaMbLE of synthetic polyploid strains. The difference in cell density between SCRaMbLE’d and non-SCRaMbLE’d synthetic and wild-type cells of each strain were plotted for the haploid (A), diploid (C) and tetraploid (E) strains. The growth profiles of haploid (B), diploid (D) and tetraploid (F) strains in the presence of 1uM estradiol. Data points and errors bars represent mean and standard deviation from three biological replicates.

## Key resources table

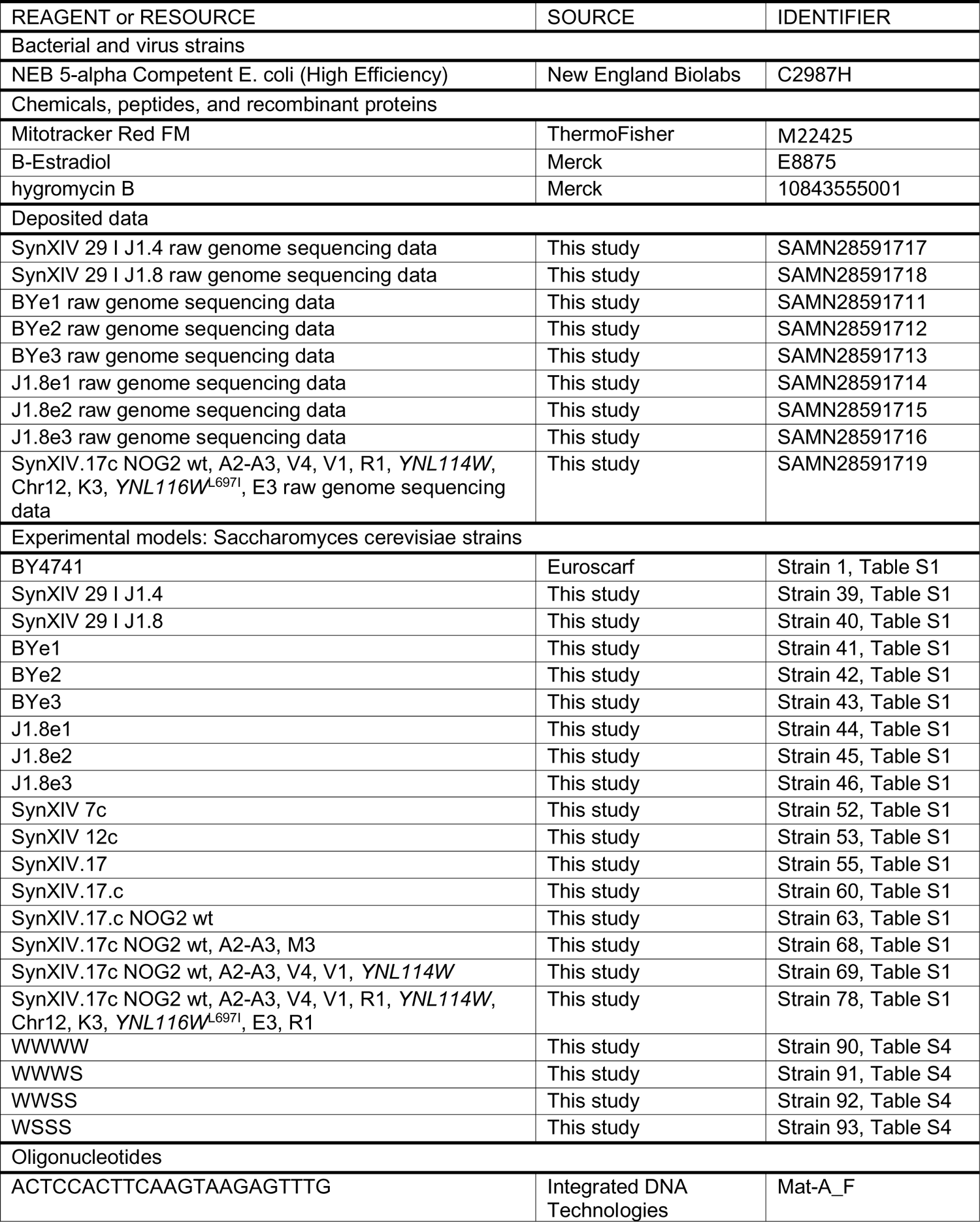
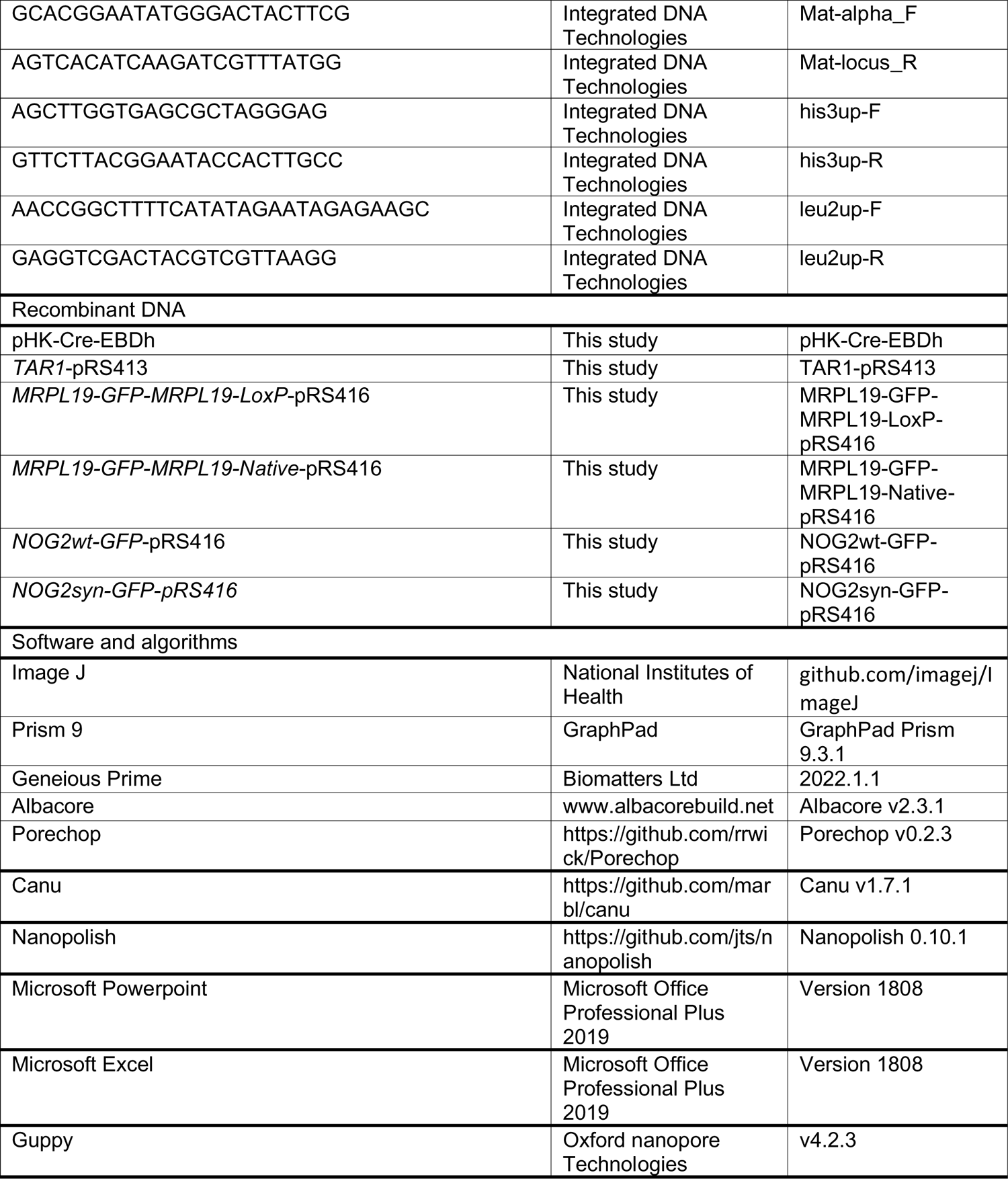

---

**Table S1.**
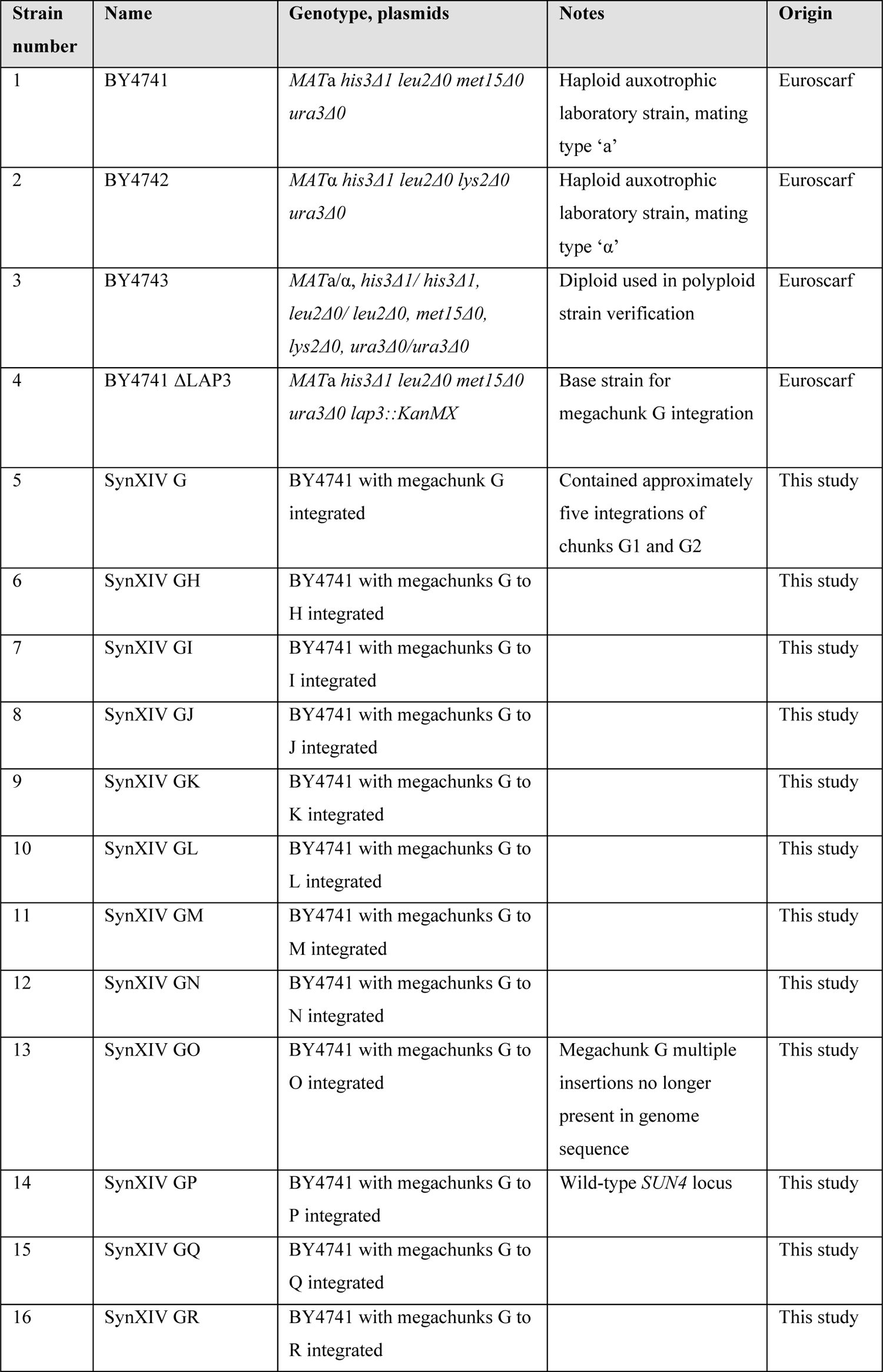
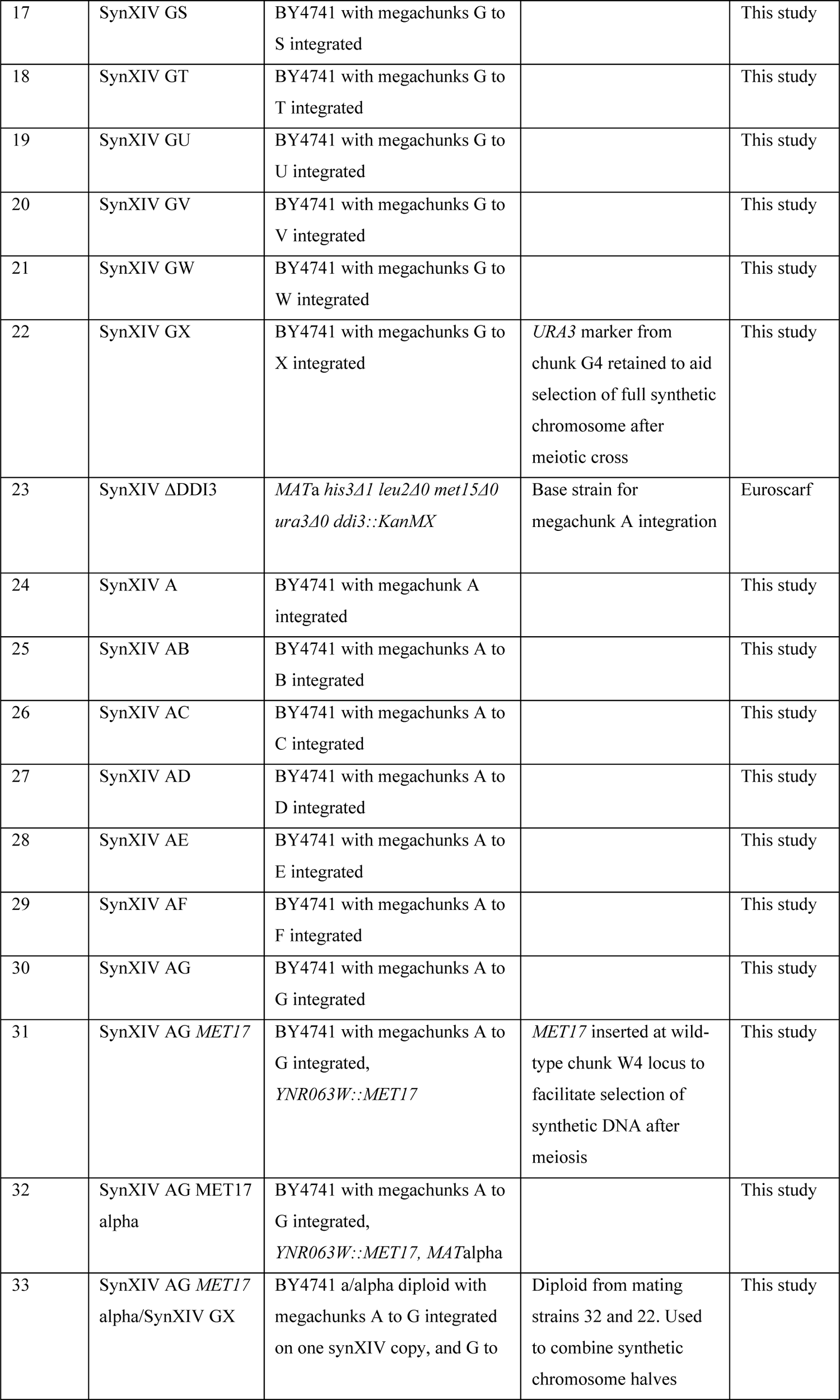
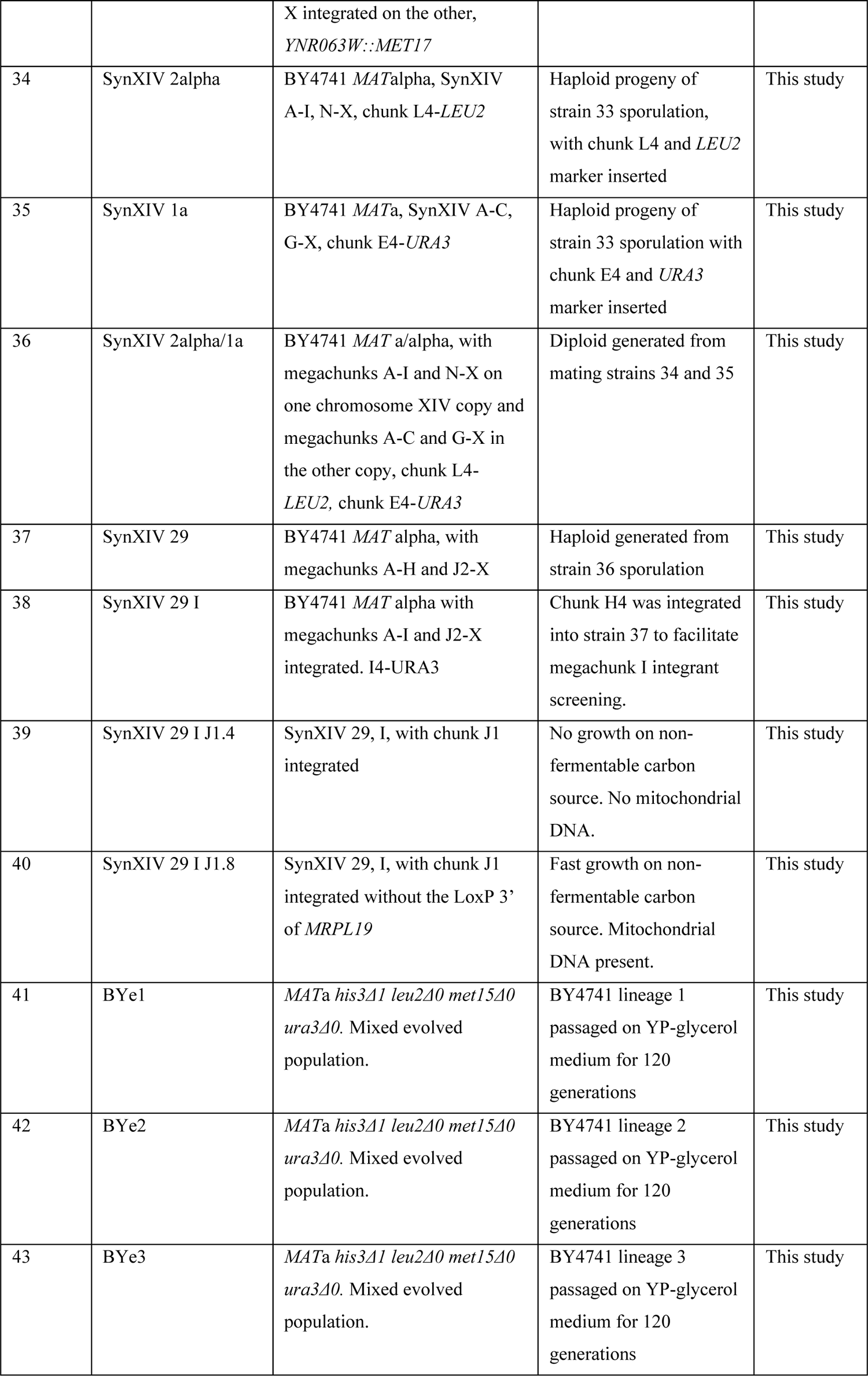
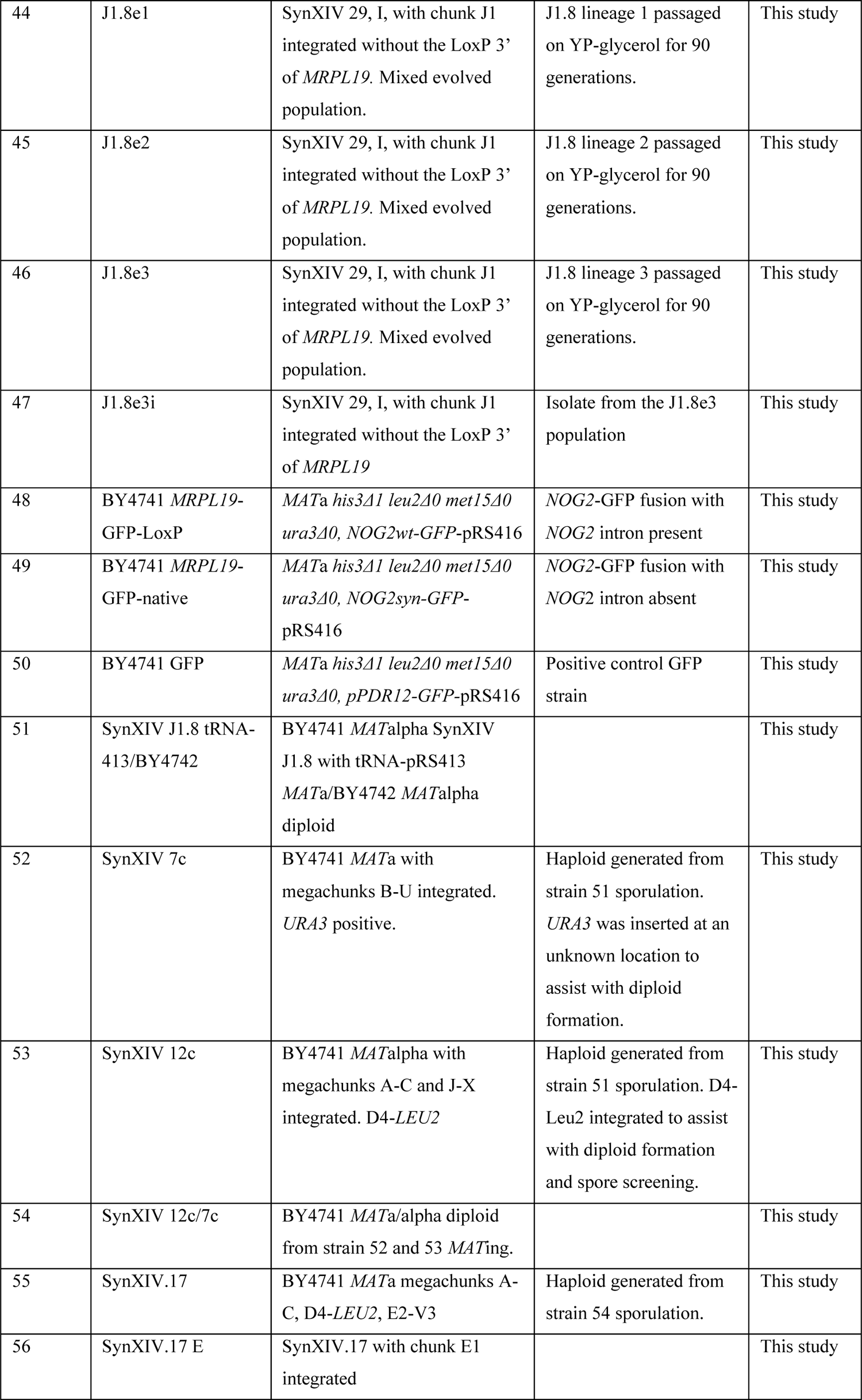
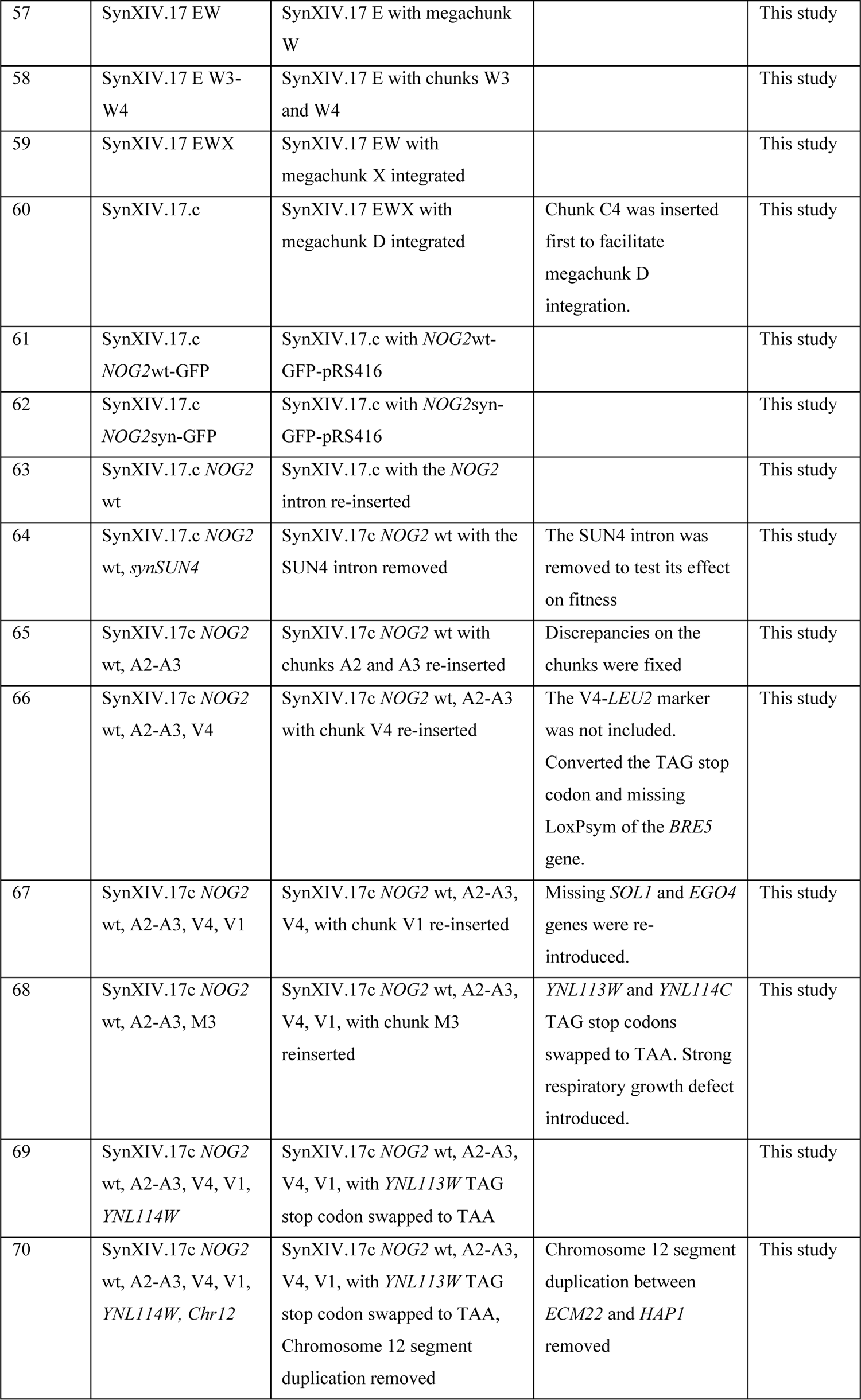
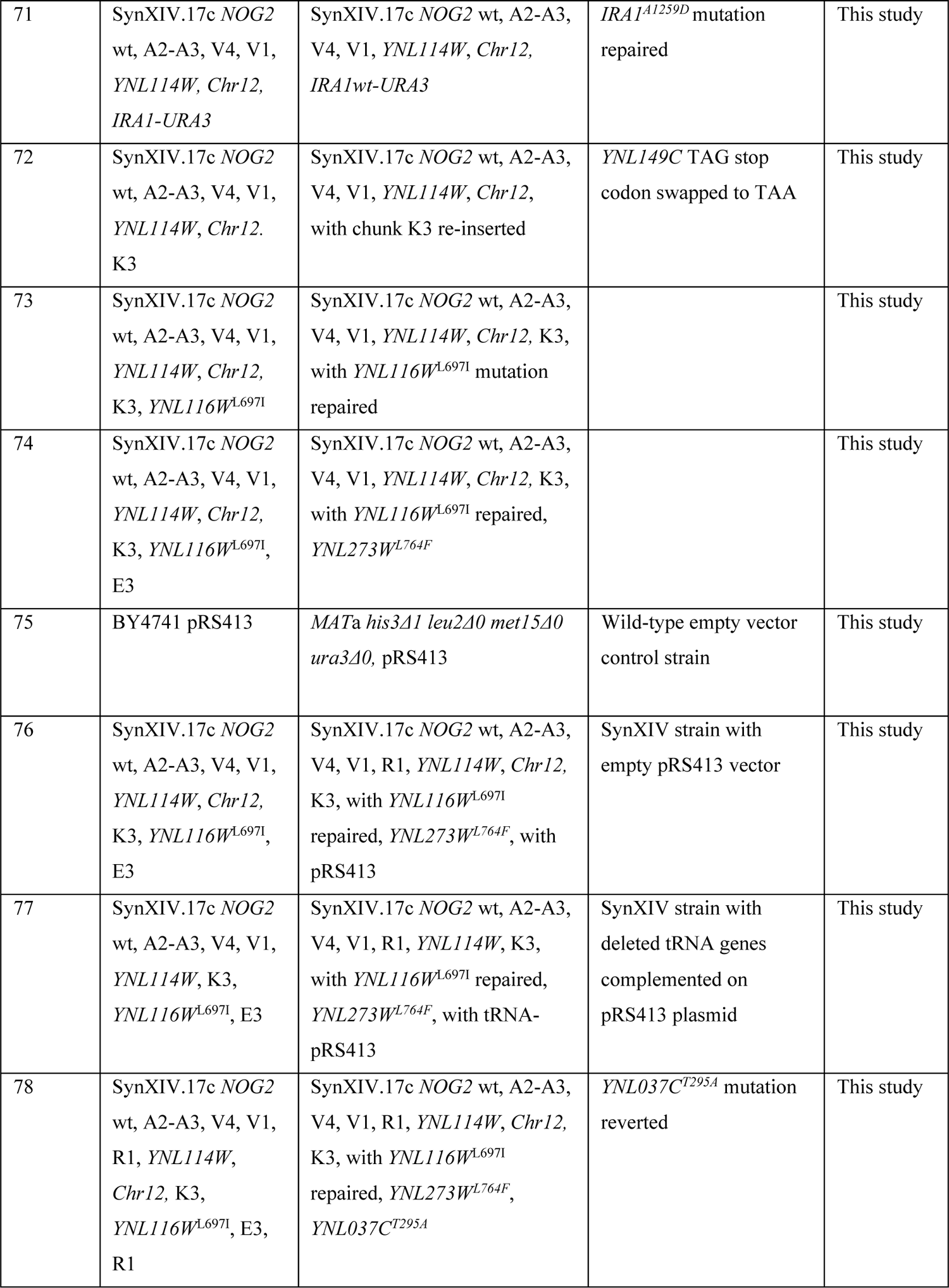
Yeast strains used in this study to build and test synXIV.

**Table S2.** SynXIV strain corrected sequence discrepancies. Whole genome sequencing of synXIV strain revealed a number of missing Sc2.0 features and point mutations that deviated from the intended synXIV sequence. A subset of these were selected for repair according to their relative importance to the project, and ease of re-introduction. All missing TAA stop codons were repaired, as this modification will serve to free up the TAA codon to encode for non-natural amino acids in the future. All non-synonymous mutations in open reading frames were repaired to enable functional expression of the relevant proteins. Some missing LoxP sites were corrected if they were nearby other features already being repaired, but were otherwise left as-is due to the fact that SCRaMbLE has a high degree of redundancy. PCR-tags are only used to verify the correct insertion of megachunks during the construction phase and were therefore left unaltered if missing, unless they were nearby another fix. Synonymous point mutations in open reading frames were also left unaltered.

| **Discrepancy Number** | **Discrepancy type** | **Original chunk/chromosome location** | **Gene(s)** | **Protein affect** |
| --- | --- | --- | --- | --- |
| 1 | T to C substitution | A3, 12735 bp | YNL329C | R276G |
| 2 | T insertion | A3, 13932 bp | YNL328C | Frame-shift |
| 3 | T to G substitution | A3, 13944 bp | YNL328C | N98H |
| 4 | C to T substitution | A3, 18438 bp | YNL326C | R302Q |
| 5 | G to T substitution | A3, 18512 bp | YNL326C | F277L |
| 6 | A to G substitution | A3, 18709 bp | YNL326C | Y212H |
| 7 | T insertion | A4, 19902 bp | YNL325C | Frame-shift |
| 8 | A to G substitution | A4, 22042 bp | YNL325C | synonymous |
| 9 | G to A transition | A4, 25505 bp | *intergenic* | - |
| 10 | G to A substitution | A4, 26370 bp | YNL321W | synonymous |
| 11 | Missing TAA stop codon | B4, 52811 bp | YNL304W | synonymous |
| 12 | Missing LoxP site | D4, 121257 bp | YNL270C | - |
| 13 | Missing TAA stop codon and LoxP site | E1, 125772 bp | YNL268W | synonymous |
| 14 | T to A substitution | E1, 125952 bp | *intergenic* | - |
| 15 | C to T substitution | E3, 110,759 bp | YNL273W | L764F |
| 16 | T to C substitution | K3, 336, 837 bp | YNL149C | None, TAA to TAG stop codon |
| 17 | Missing TAA stop codon | M3, 400807 bp | YNL113C | synonymous |
| 18 | Missing *Bsu36I* restriction site | M3-M4, 401355 bp | *YNL112W* | synonymous |
| 19 | T to C substitution | R1, 539,674 bp | *YNL037C* | T295A |
| 20 | Chunk V1 missing | V1, 2972 bp deletion beginning at 663,572 bp |  | YNR034 and YNR034W looped out between LoxP sites |
| 21 | Missing LoxP site | V4, 690721 bp | *YNR051C* | - |
| 22 | Missing TAA stop codon | V4, 690758 bp | *YNR051C* | synonymous |
| 23 | Missing PCR-tag | V4, 690776 bp | *YNR051C* | synonymous |
| 24 | Duplication on chromosome 12 | *ECM22* to *HAP1* |  |  |

**Table S3.**
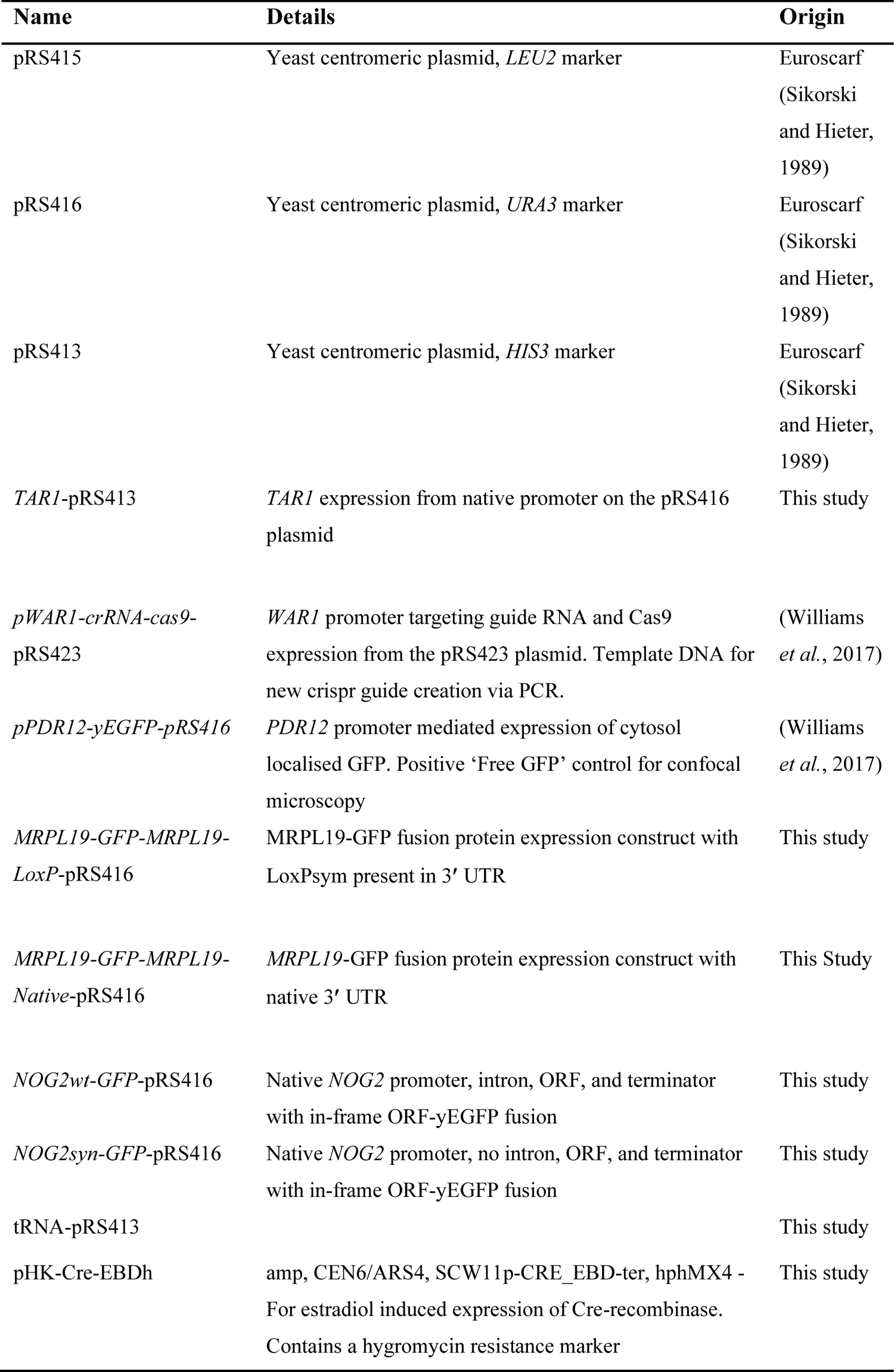
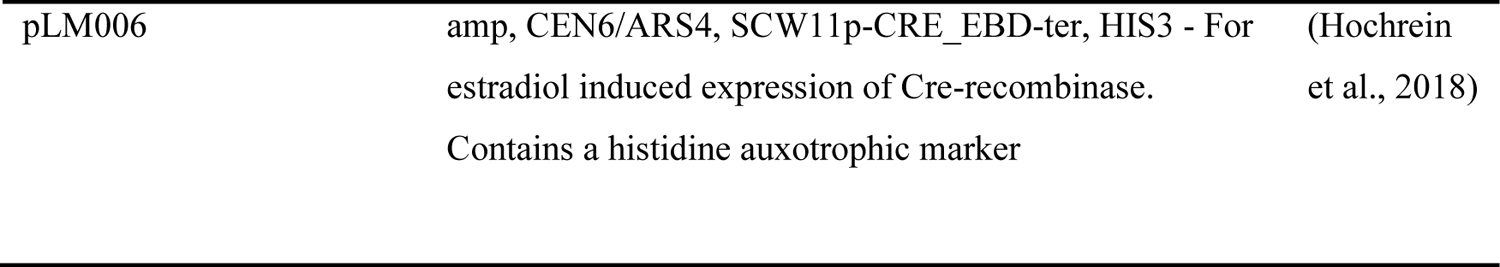
Plasmids used in this study.

| **Name** | **Details** | **Origin** |
| --- | --- | --- |
| pRS415 | Yeast centromeric plasmid, *LEU2* marker | Euroscarf<br>(Sikorski<br>and Hieter,<br>1989) |
| pRS416 | Yeast centromeric plasmid, *URA3* marker | Euroscarf<br>(Sikorski<br>and Hieter,<br>1989) |
| pRS413 | Yeast centromeric plasmid, *HIS3* marker | Euroscarf<br>(Sikorski<br>and Hieter,<br>1989) |
| *TAR1*-pRS413 | *TAR1* expression from native promoter on the pRS416 plasmid | This study |
| *pWAR1-crRNA-cas9*-<br>pRS423 | *WAR1* promoter targeting guide RNA and Cas9 expression from the pRS423 plasmid. Template DNA for new crispr guide creation via PCR. | (Williams<br>*et al.*, 2017) |
| *pPDR12-yEGFP*-pRS416 | *PDR12* promoter mediated expression of cytosol localised GFP. Positive ‘Free GFP’ control for confocal microscopy | (Williams<br>*et al.*, 2017) |
| *MRPL19-GFP-MRPL19*-<br>*LoxP*-pRS416 | *MRPL19*-GFP fusion protein expression construct with *LoxP*sym present in 3’ UTR | This study |
| *MRPL19-GFP-MRPL19*-<br>*Native*-pRS416 | *MRPL19*-GFP fusion protein expression construct with native 3’ UTR | This Study |
| *NOG2wt-GFP*-pRS416 | Native *NOG2* promoter, intron, ORF, and terminator with in-frame ORF-yEGFP fusion | This study |
| *NOG2syn-GFP*-pRS416 | Native *NOG2* promoter, no intron, ORF, and terminator with in-frame ORF-yEGFP fusion | This study |
| tRNA-pRS413 |  | This study |
| pHK-Cre-EBDh | amp, CEN6/ARS4, SCW11p-CRE_EBD-ter, hphMX4 - For estradiol induced expression of Cre-recombinase. Contains a hygromycin resistance marker | This study |
| pLM006 | amp, CEN6/ARS4, SCW11p-CRE_EBD-ter, HIS3 - For<br>estradiol induced expression of Cre-recombinase.<br>Contains a histidine auxotrophic marker | (Hochrein<br>et al., 2018) |

**Table S4.**
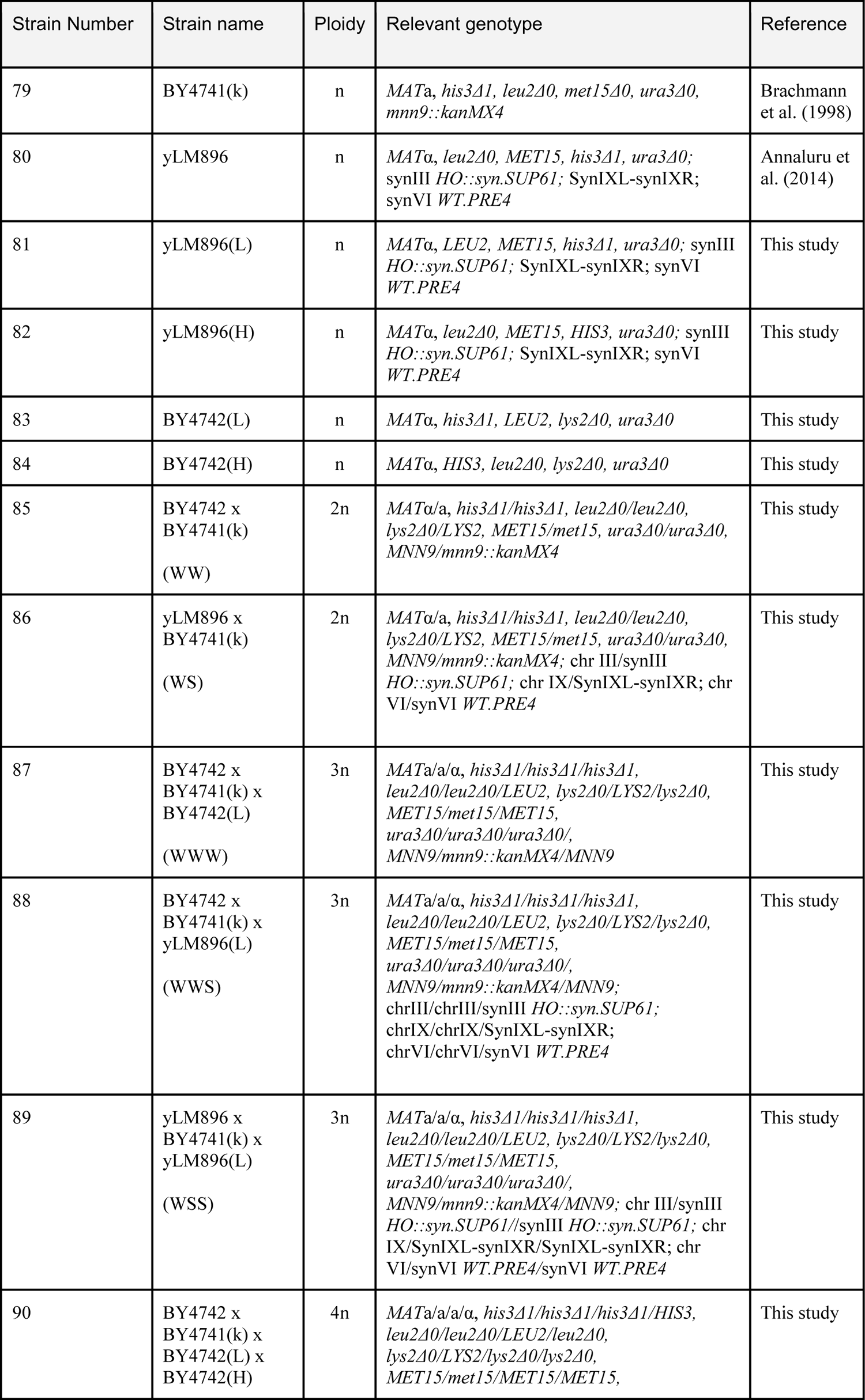
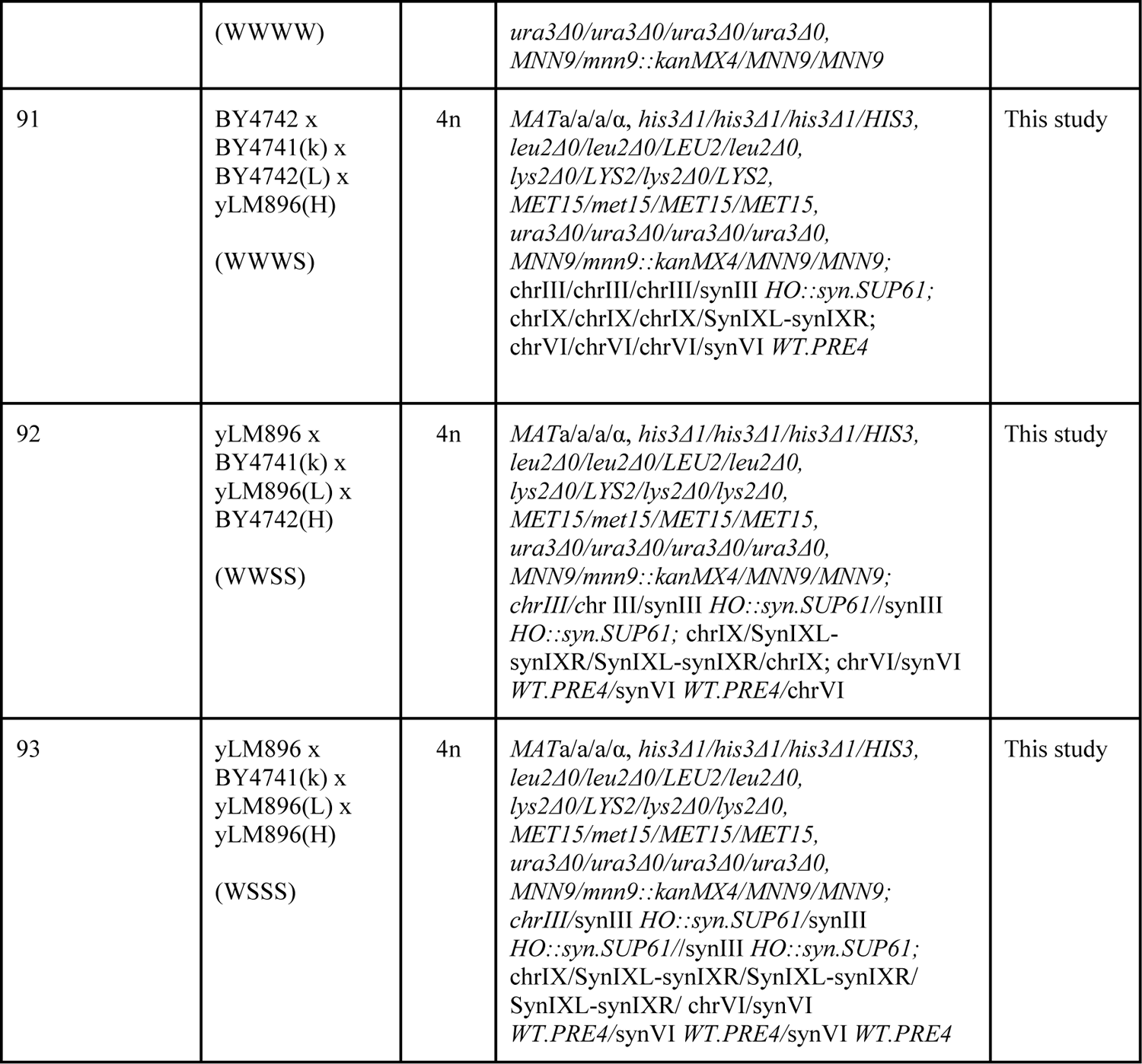
Yeast strains used for polyploid construction.

| Strain Number | Strain name | Ploidy | Relevant genotype | Reference |
| --- | --- | --- | --- | --- |
| 79 | BY4741(k) | n | *MATa, his3Δ1, leu2Δ0, met15Δ0, ura3Δ0, mnn9::kanMX4* | Brachmann et al. (1998) |
| 80 | yLM896 | n | *MATα, leu2Δ0, MET15, his3Δ1, ura3Δ0; synIII HO::syn.SUP61; SynIXL-synIXR; synVI WT.PRE4* | Annaluru et al. (2014) |
| 81 | yLM896(L) | n | *MATα, LEU2, MET15, his3Δ1, ura3Δ0; synIII HO::syn.SUP61; SynIXL-synIXR; synVI WT.PRE4* | This study |
| 82 | yLM896(H) | n | *MATα, leu2Δ0, MET15, HIS3, ura3Δ0; synIII HO::syn.SUP61; SynIXL-synIXR; synVI WT.PRE4* | This study |
| 83 | BY4742(L) | n | *MATα, his3Δ1, LEU2, lys2Δ0, ura3Δ0* | This study |
| 84 | BY4742(H) | n | *MATα, HIS3, leu2Δ0, lys2Δ0, ura3Δ0* | This study |
| 85 | BY4742 x<br>BY4741(k)<br>(WW) | 2n | *MATα/a, his3Δ1/his3Δ1, leu2Δ0/leu2Δ0, lys2Δ0/LYS2, MET15/met15, ura3Δ0/ura3Δ0, MNN9/mnn9::kanMX4* | This study |
| 86 | yLM896 x<br>BY4741(k)<br>(WS) | 2n | *MATα/a, his3Δ1/his3Δ1, leu2Δ0/leu2Δ0, lys2Δ0/LYS2, MET15/met15, ura3Δ0/ura3Δ0, MNN9/mnn9::kanMX4; chr III/synIII HO::syn.SUP61; chr IX/SynIXL-synIXR; chr VI/synVI WT.PRE4* | This study |
| 87 | BY4742 x<br>BY4741(k) x<br>BY4742(L)<br>(WWW) | 3n | *MATα/a/α, his3Δ1/his3Δ1/his3Δ1, leu2Δ0/leu2Δ0/LEU2, lys2Δ0/LYS2/lys2Δ0, MET15/met15/MET15, ura3Δ0/ura3Δ0/ura3Δ0/, MNN9/mnn9::kanMX4/MNN9* | This study |
| 88 | BY4742 x<br>BY4741(k) x<br>yLM896(L)<br>(WWS) | 3n | *MATα/a/α, his3Δ1/his3Δ1/his3Δ1, leu2Δ0/leu2Δ0/LEU2, lys2Δ0/LYS2/lys2Δ0, MET15/met15/MET15, ura3Δ0/ura3Δ0/ura3Δ0/, MNN9/mnn9::kanMX4/MNN9; chrIII/chrIII/synIII HO::syn.SUP61; chrIX/chrIX/SynIXL-synIXR; chrVI/chrVI/synVI WT.PRE4* | This study |
| 89 | yLM896 x<br>BY4741(k) x<br>yLM896(L)<br>(WSS) | 3n | *MATα/a/α, his3Δ1/his3Δ1/his3Δ1, leu2Δ0/leu2Δ0/LEU2, lys2Δ0/LYS2/lys2Δ0, MET15/met15/MET15, ura3Δ0/ura3Δ0/ura3Δ0/, MNN9/mnn9::kanMX4/MNN9; chr III/synIII HO::syn.SUP61/synIII HO::syn.SUP61; chr IX/SynIXL-synIXR/SynIXL-synIXR; chr VI/synVI WT.PRE4/synVI WT.PRE4* | This study |
| 90 | BY4742 x<br>BY4741(k) x<br>BY4742(L) x<br>BY4742(H) | 4n | *MATα/a/a/α, his3Δ1/his3Δ1/his3Δ1/HIS3, leu2Δ0/leu2Δ0/LEU2/leu2Δ0, lys2Δ0/LYS2/lys2Δ0/lys2Δ0, MET15/met15/MET15/MET15,* | This study |
|  | (WWWW) |  | *ura3Δ0/ura3Δ0/ura3Δ0/ura3Δ0, MNN9/mnn9::kanMX4/MNN9/MNN9* |  |
| 91 | BY4742 x<br>BY4741(k) x<br>BY4742(L) x<br>yLM896(H)<br>(WWWS) | 4n | *MATa/a/a/α, his3Δ1/his3Δ1/his3Δ1/HIS3, leu2Δ0/leu2Δ0/LEU2/leu2Δ0, lys2Δ0/LYS2/lys2Δ0/LYS2, MET15/met15/MET15/MET15, ura3Δ0/ura3Δ0/ura3Δ0/ura3Δ0, MNN9/mnn9::kanMX4/MNN9/MNN9; chrIII/chrIII/chrIII/synIII HO::syn.SUP61; chrIX/chrIX/chrIX/SynIXL-synIXR; chrVI/chrVI/chrVI/synVI WT.PRE4* | This study |
| 92 | yLM896 x<br>BY4741(k) x<br>yLM896(L) x<br>BY4742(H)<br>(WWSS) | 4n | *MATa/a/a/α, his3Δ1/his3Δ1/his3Δ1/HIS3, leu2Δ0/leu2Δ0/LEU2/leu2Δ0, lys2Δ0/LYS2/lys2Δ0/lys2Δ0, MET15/met15/MET15/MET15, ura3Δ0/ura3Δ0/ura3Δ0/ura3Δ0, MNN9/mnn9::kanMX4/MNN9/MNN9; chrIII/chr III/synIII HO::syn.SUP61//synIII HO::syn.SUP61; chrIX/SynIXL-synIXR/SynIXL-synIXR/chrIX; chrVI/synVI WT.PRE4/synVI WT.PRE4/chrVI* | This study |
| 93 | yLM896 x<br>BY4741(k) x<br>yLM896(L) x<br>yLM896(H)<br>(WSSS) | 4n | *MATa/a/a/α, his3Δ1/his3Δ1/his3Δ1/HIS3, leu2Δ0/leu2Δ0/LEU2/leu2Δ0, lys2Δ0/LYS2/lys2Δ0/lys2Δ0, MET15/met15/MET15/MET15, ura3Δ0/ura3Δ0/ura3Δ0/ura3Δ0, MNN9/mnn9::kanMX4/MNN9/MNN9; chrIII/synIII HO::syn.SUP61/synIII HO::syn.SUP61//synIII HO::syn.SUP61; chrIX/SynIXL-synIXR/SynIXL-synIXR/ SynIXL-synIXR/ chrVI/synVI WT.PRE4/synVI WT.PRE4/synVI WT.PRE4* | This study |

**Table S5.** Strain version table.

| Version name | Strain number | Comment | Details |
| --- | --- | --- | --- |
| yeast_chr14_0_00 | NA | Wild-type chromosome 14 sequence | GenBank: BK006947.3 |
| yeast_chr14_3_26 | NA | Original design sequence | Final design by BioStudio |
| yeast_chr14_9_01 | SynXIV 29 I J1.4, Strain 39, Table S1 | synXIV Draft strain, with 5 TAG stop codons, 10 loxPsym sites missing, 23 wild-type, 4 point mutations causing amino acid changes, *YNL066W* intron present. | **Remaining TAG stop codons:** 125772 A->G, 336549 T->C, 400292 T->C, 400807 A->G, 690758 T->C. **Missing loxPsym sites:** 120565-120598, 125051-125084, 250019-250052, 337131-337164, 374188-374221, 438943-438976, 603336-603369, 690721-690754, 746919-746952, 747387-747420. **Wild-type PCRTags:** 108692-109106 YNL273W_525, 131105-131132 YNL264C_193,174923-174950 YNL243W_1380, 175389-175416 YNL243W_2607, 180372-180399 YNL242W_3522, 249297-249324 YNL201C_2362, 250194-250221 YNL200C_139, 250428-250455 YNL200C_373, 281076-281103 YNL183C_1189, 281403-281430 YNL183C_1516, 337177-337204 YNL148C_10, 337465-337492 YNL148C_298, 341073-341100 YNL144C_1021, 410750-410777 YNL106C_3133, 489097-489118 YNL066W_673, 491634-491661 YNL065W_609, 491925-491952 YNL065W_900, 495487-495514 YNL063W_360, 595657-595684 YNL005C_388, 658820-658847 YNR031C_4591, 689809-689836 YNR051C_19, 725470-725497 YNR065C_2761, 747793-747820 YNR073C_370, 748138-748165 YNR073C_715. **Point mutations that cause amino acid changes:** 12071 G->A, 140826 C->T, 396611 T->A, 450624 T->C. |
| yeast_chr14_9_02 | SynXIV 29 I J1.8, Strain 40, Table S1 | synXIV Draft strain, removed *MRPL19* LoxP site | **Remaining TAG stop codons:** 125772 A->G, 336549 T->C, 400292 T->C, 400807 A->G, 690758 T->C. **Missing loxPsym sites:** 120565-120598, 125051-125084, 250019-250052, 337131-337164, 374188-374221, 438943-438976, 603336-603369, 690721-690754, 746919-746952, 747387-747420, 278873-278906. **Wild-type PCRTags:** 108692-109106 YNL273W_525, 131105-131132 YNL264C_193,174923-174950 YNL243W_1380, 175389-175416 YNL243W_2607, 180372-180399 YNL242W_3522, 249297-249324 YNL201C_2362, 250194-250221 YNL200C_139, 250428-250455 YNL200C_373, 281076-281103 YNL183C_1189, 281403-281430 YNL183C_1516, 337177-337204 YNL148C_10, 337465-337492 YNL148C_298, 341073-341100 YNL144C_1021, 410750-410777 YNL106C_3133, 489097-489118 YNL066W_673, 491634-491661 YNL065W_609, 491925-491952 YNL065W_900, 495487-495514 YNL063W_360, 595657-595684 YNL005C_388, 658820-658847 YNR031C_4591, 689809-689836 YNR051C_19, 725470-725497 YNR065C_2761, 747793-747820 YNR073C_370, 748138-748165 YNR073C_715. **Point mutations that cause amino acid changes:** 12071 G->A, 140826 C->T, 396611 T->A, 450624 T->C. |
| yeast_chr14_9_03 | SynXIV.17c NOG2 wt, A2-A3, V4, V1, R1, *YNL114W*, | SynXIV draft strain. 3 TAG stop codons changed to TAA, 4 LoxPsym sites added, 4 | **Remaining TAG stop codons:** 400292 T->C. **Missing loxPsym sites:** 125051-125084, 250019-250052, 278873-278906, 337131-337164, 374188-374221, 438943-438976, 603336-603369, 715110-715143. **Wild-type PCRTags:** 108692-109106 YNL273W_525,116368-116395 YNL271C_1528, 124292-124319 YNL268W_390, |
|  | Chr12, K3, *YNL116W*<sup>L697I</sup>, E3. Strain 77, Table S1 | non-synonymous point mutations corrected, 3 PCR-tags added, and 18 PCR tags removed during reconstruction and repair. *NOG2* intron re-inserted, | 124640-124667 *YNL268W*_738, 124700-124727 *YNL268W*_798, 124901-124928 *YNL268W*_999, 126914-126941 *YNL267W*_684, 127169-127196 *YNL267W*_939, 128195-128222 *YNL267W*_1965, 128627-128654 *YNL267W*_2397, 129047-129074 *YNL267W*_2817, 129242-129269 *YNL267W*_3012, 131105-131132 *YNL264C*_193, 174923-174950 *YNL243W*_1380, 175389-175416 *YNL243W*_2607, 180372-180399 *YNL242W*_3522, 249297-249324 *YNL201C*_2362, 250194-250221 *YNL200C*_139, 250428-250455 *YNL200C*_373, 337177-337204 *YNL148C*_10, 337465-337492 *YNL148C*_298, 341073-341100 *YNL144C*_1021, 410750-410777 *YNL106C*_3133, 489097-489118 *YNL066W*_673, 491634-491661 *YNL065W*_609, 491925-491952 *YNL065W*_900, 495487-495514 *YNL063W*_360, 595657-595684 *YNL005C*_388, 658820-658847 *YNR031C*_4591, 691283-691310 *YNR051C*_484, 691337-691364 *YNR051C*_538, 691670-691697 *YNR051C*_871, 696935-696962 *YNR054C*_58, 697238-697265 *YNR054C*_361, 715318-715345 *YNR061C*_172, 715507-715534 *YNR061C*_361. **Point mutations that cause amino acid changes:** 450624 T->C. |
| yeast_chr14_9_04 | SynXIV.17c *NOG2* wt, A2-A3, V4, V1, R1, *YNL114W*, Chr12, K3, *YNL116W*<sup>L697I</sup>, E3, R1. Strain 78, Table S1 | SynXIV final strain. Non-synonymous point mutation in *IDH1* reverted to wild-type sequence | **Remaining TAG stop codons:** 400292 T->C. **Missing loxPsym sites:** 125051-125084, 250019-250052, 278873-278906, 337131-337164, 374188-374221, 438943-438976, 603336-603369, 715110-715143. **Wild-type PCRTags:** 108692-109106 *YNL273W*_525, 116368-116395 *YNL271C*_1528, 124292-124319 *YNL268W*_390, 124640-124667 *YNL268W*_738, 124700-124727 *YNL268W*_798, 124901-124928 *YNL268W*_999, 126914-126941 *YNL267W*_684, 127169-127196 *YNL267W*_939, 128195-128222 *YNL267W*_1965, 128627-128654 *YNL267W*_2397, 129047-129074 *YNL267W*_2817, 129242-129269 *YNL267W*_3012, 131105-131132 *YNL264C*_193, 174923-174950 *YNL243W*_1380, 175389-175416 *YNL243W*_2607, 180372-180399 *YNL242W*_3522, 249297-249324 *YNL201C*_2362, 250194-250221 *YNL200C*_139, 250428-250455 *YNL200C*_373, , 337177-337204 *YNL148C*_10, 337465-337492 *YNL148C*_298, 341073-341100 *YNL144C*_1021, 410750-410777 *YNL106C*_3133, 489097-489118 *YNL066W*_673, 491634-491661 *YNL065W*_609, 491925-491952 *YNL065W*_900, 495487-495514 *YNL063W*_360, 595657-595684 *YNL005C*_388, 658820-658847 *YNR031C*_4591, 691283-691310 *YNR051C*_484, 691337-691364 *YNR051C*_538, 691670-691697 *YNR051C*_871, 696935-696962 *YNR054C*_58, 697238-697265 *YNR054C*_361, 715318-715345 *YNR061C*_172, 715507-715534 *YNR061C*_361. |

